# Evolutionary and acclimatory shifts in ion transporter gene expression across salinity transitions in the copepod *Eurytemora carolleeae*

**DOI:** 10.64898/2026.11.03.763763

**Authors:** Teresa E Popp, Eva Blondeau-Bidet, Antoine Fraimout, Delphine Bonnet, Romain Gros, Carol Eunmi Lee, Catherine Lorin-Nebel

## Abstract

Many aquatic invaders are crossing biogeographic boundaries from saline to freshwater habitats. This transition often requires acclimation or evolution of physiological mechanisms to maintain internal ionic homeostasis. Populations of the copepod *Eurytemora affinis* complex have repeatedly invaded freshwater environments from saline habitats over the past century, providing an ideal model for investigating rapid adaptation to salinity change. Prior studies have found that ion transport-related genes form the largest functional categories of genes under selection and show evolutionary shifts in gene expression across saline to freshwater transitions in the *E. affinis* complex. In this study, focusing on the top targets of selection, we investigated evolutionary and acclimatory shifts in gene expression for 19 ion transport-related genes for populations of *E. carolleeae* (Atlantic clade of the *E. affinis* complex). Using a quantitative real-time PCR approach, we compared gene expression between ancestral saline and derived freshwater populations through a common garden reaction norm experiment and a laboratory natural selection experiment. We found that different ion transport-related gene paralogs within expanded gene families showed divergent patterns of expression in response to salinity change, consistent with specialization. For instance, Na^+^/K^+^-ATPase subunit *α* paralog 1 (*NKA-α-1)* and Na^+^,K^+^,2Cl^-^ Cotransporter paralog 3 *(NKCC-3)* exhibited the evolution of increased expression in the freshwater population across all conditions, relative to the saline population. In contrast, *Na^+^/H^+^ Antiporter* paralog 7 *(NHA-7)* and *NHA-5* showed evolution of elevated expression in the saline population under low salinity relative to the freshwater population. Several genes also showed evolutionary differences in plasticity between saline and freshwater populations, such that saline populations retained phenotypic plasticity, whereas freshwater populations were comparatively canalized. In the laboratory natural selection experiment imposing salinity decline, evolutionary responses of the paralogs *NKCC-3*, *NHA-5*, and *NHA-7* in the saline selection line largely recapitulated the evolutionary shifts observed in the natural populations. On the other hand, expression patterns of the ion transporters in the freshwater selection line showed little response during salinity increase, suggesting canalization. Together, these results reveal evolutionary and acclimatory divergence in gene expression among paralogs, suggesting the evolution of functional divergence among them.

**Graphical Abstract:** 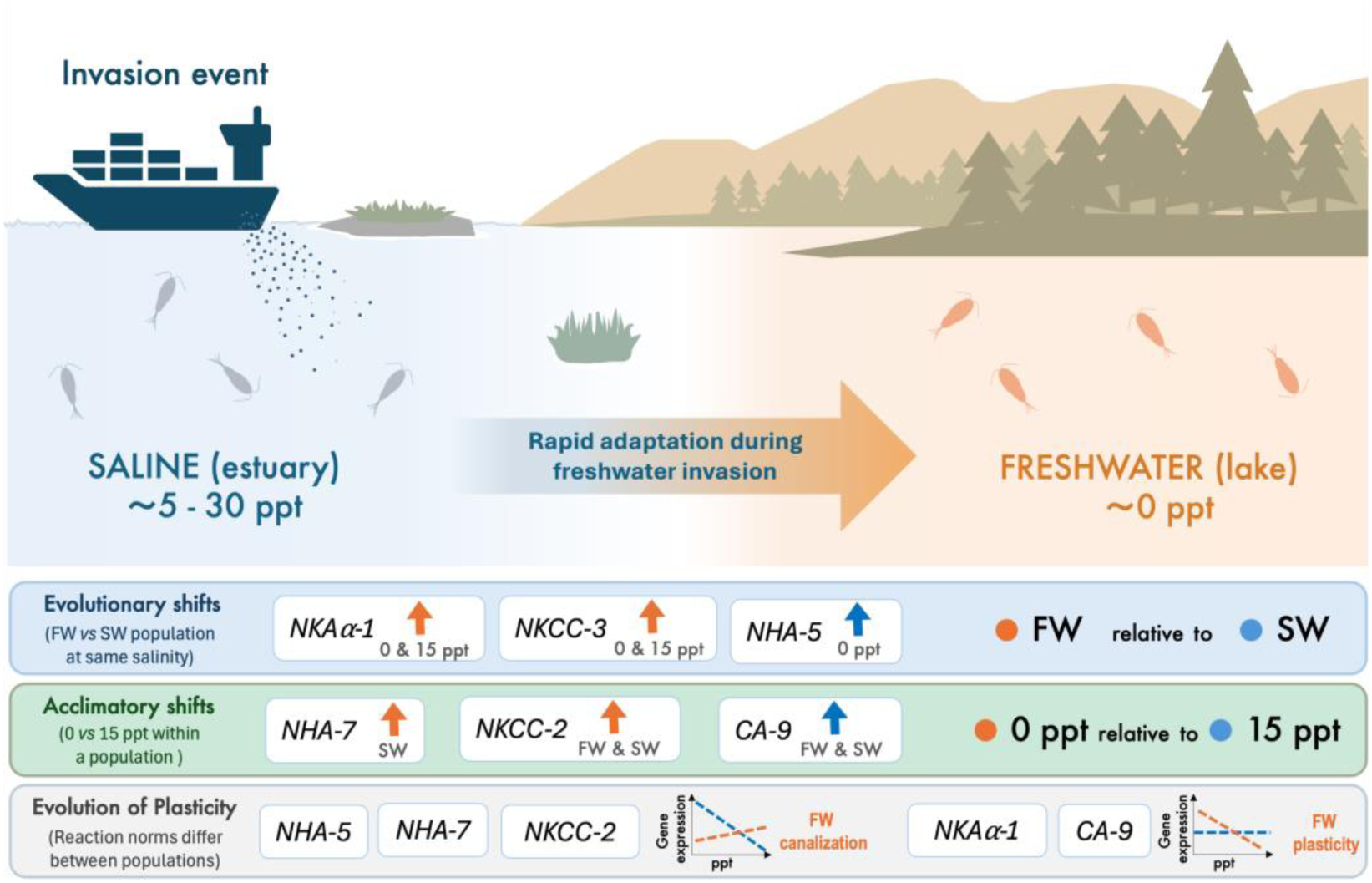

## INTRODUCTION

Invasive populations often encounter rapid and drastic environmental changes following introductions into novel habitats. Such rapid changes include abiotic and biotic factors, such as temperature, salinity, competition, predation, and food availability (1–3). Consequently, only a small fraction of introduced species survive the novel conditions and successfully become established (4,5). Determining the ability of organisms to acclimate or populations to evolve in response to novel conditions is crucial for understanding their capacity to survive and avoid extinction under changing environments (5–7).

Notably, freshwater environments are increasingly being dominated by species invading from saline habitats (8–10). For example, the Great Lakes of North America have experienced a large influx of invaders from the brackishwater Black and Caspian (Ponto-Caspian) Sea region (8,10–12). Brackishwater species, such as the quagga mussel *Dreissena bugensis*, the zebra mussel *Dreissena polymorpha*, and the fishhook water flea *Cercopagis pengoi* (13–16), were likely introduced *via* cargo ship ballast water following the opening of the Saint Lawrence Seaway in 1959 (11,12). However, this transition between saline and freshwater habitats (particularly around the 5 ppt boundary) represents a formidable biogeographic barrier that poses serious physiological challenges for organisms in maintaining internal ionic homeostasis (17–20). Ultimately, ion transporters (see details below) likely play pivotal roles in enabling organisms and populations to acclimate and adapt to changes in salinity (21–23).

Evolution and acclimation of physiological processes often play crucial roles in the success of invasive populations across various stages of invasion, including transport, introduction, establishment, and range expansion (6,7,24). The extent of phenotypic plasticity of organisms and the evolvability of populations ultimately shapes their responses to rapid environmental change. Individual acclimatory responses (resulting from phenotypic plasticity) are often critically important for the initial stages of survival in a novel environment. However, when the extent of environmental change exceeds the physiological tolerance and/or limits of individual plastic responses, rapid evolutionary changes are often necessary for survival of the population in the novel environment (6,7). While many studies have examined plastic responses of ion transporter expression or function across different salinities (25–28), relatively few studies have examined evolutionary responses of these ion transporters to changes in salinity (23,27,29–31).

The copepod *Eurytemora affinis* species complex provides a valuable model system for investigating physiological mechanisms of adaptation and acclimation in response to salinity change (32,33). Estuarine populations of the *E. affinis* complex have independently colonized freshwater habitats across the Northern Hemisphere over the past ∼80 years, likely due to the transport and dumping of ship ballast water (11,12,34,35) (Fig 1). By investigating these recent freshwater invasions, we can gain insights into how populations might respond to future changes in habitat salinity induced by climate change (3).

**Fig 1.**
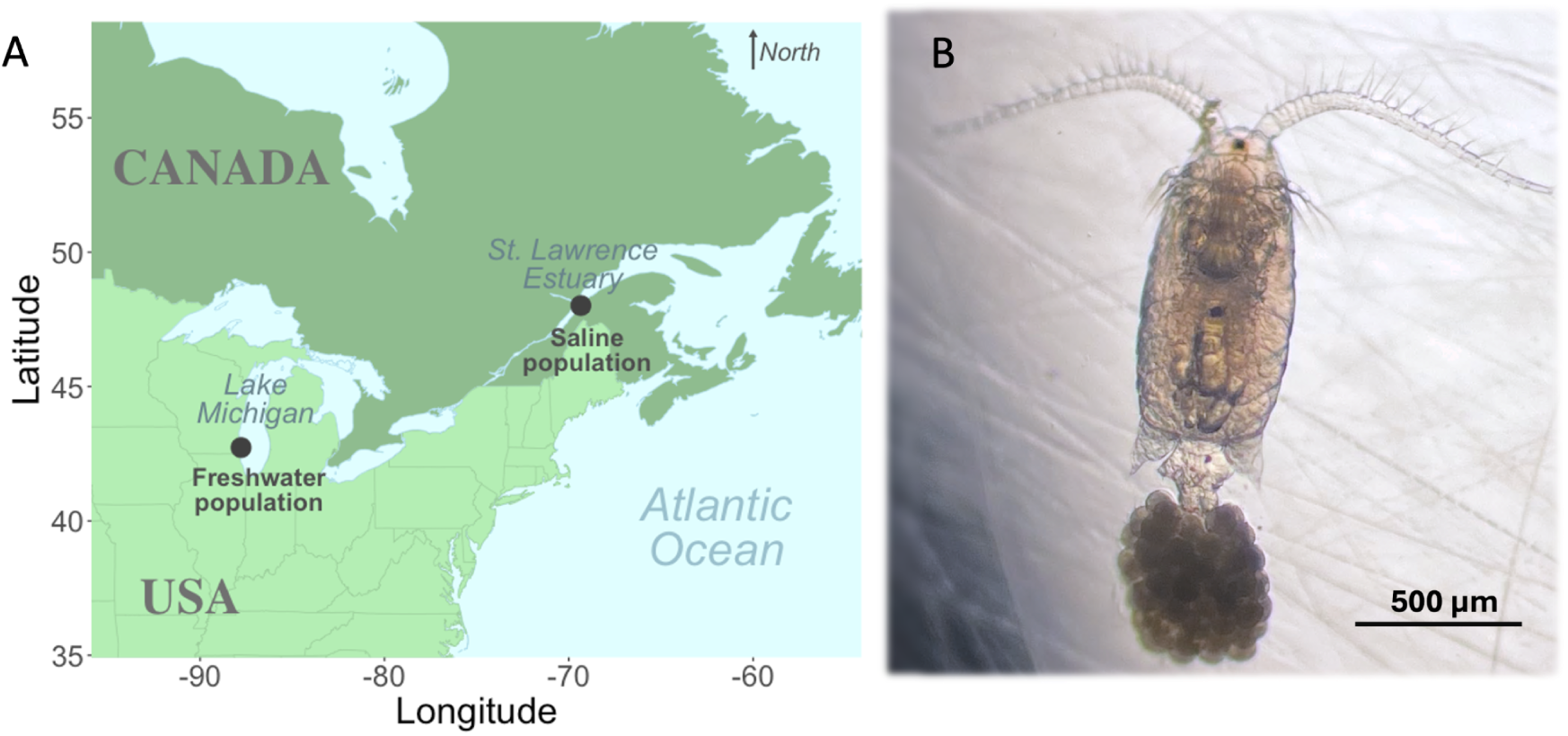
Populations of *Eurytemora carolleeae* (Atlantic clade of the *E. affinis* complex) used in this study. **(A)** Black dots indicate the locations of the saline and freshwater populations used in this study. The ancestral saline population in Baie de L’Isle Verte (in the St. Lawrence estuary) experiences seasonally fluctuating salinities from 5 to 40 ppt, whereas the invasive freshwater population in Lake Michigan resides in a relatively constant salinity of 0 ppt (conductivity of ∼300 *μ*S/cm). **(B)** Dorsal view of an ovigerous adult female from the freshwater population in Lake Michigan. Photo by Teresa E. Popp and Catherine Lorin-Nebel.

Prior studies on the evolutionary adaptation of *E. affinis* complex populations during salinity transitions have identified genes involved in ion transport as major targets of natural selection in response to salinity change (23,29,36–38). Specifically, a set of ion transport-related gene paralogs repeatedly showed signatures of selection (allele frequency shifts) between saline and freshwater populations. These include paralogs within gene families of *Na^+^/H^+^ Antiporter* (*NHA*), *Na^+^/K^+^-ATPase* (*NKA*), *Na^+^,K^+^,2Cl^-^ Cotransporter* (*NKCC*), *Carbonic Anhydrase* (*CA*), and *V-type H^+^ ATPase* (*VHA*) (36–40). Many of these genes that displayed signatures of selection also showed evolutionary shifts in gene expression based on Illumina RNA-seq between saline and freshwater populations of *E. carolleeae* (Atlantic clade of the *E. affinis* complex) (29). In addition, *in situ* immunolocalization studies further revealed that several of these ion transporter proteins were expressed in ionocytes (41,42), which are specialized cells involved in ion transport and the regulation of ion homeostasis. In *E. affinis* complex, these cells occur in putative osmoregulatory organs, including in the swimming legs, maxillary glands, and midgut (30,31,41,42). Moreover, within these organs, the ion transporters NHA and NKA showed evolutionary shifts in *in situ* expression between saline and freshwater populations, as well as acclimatory shifts across salinities (30,31). Together, these results suggest that these ion transport-related paralogs contribute to freshwater acclimation and adaptation.

Notably, variation in patterns of gene expression and signatures of selection among individual ion transport-related paralogs suggest that they are potentially not redundant in function and contribute differentially to salinity adaptation. Variation in tissue localization, gene and protein expression patterns, and signatures of selection among the ion transport-related paralogs (21,29,31,43) suggest functional divergence among them. Specifically, individual paralogs of the ion transport-related gene families vary in the direction of gene expression in response to salinity change, in terms of both evolutionary shifts between saline and freshwater populations and acclimatory shifts across salinities (21,29). In addition, distinct NHA paralogs show variation in patterns of localization and differences in acclimatory (between salinities) and evolutionary (between populations) shifts in protein expression within osmoregulatory organs (31).

In addition, potential functional divergence among the ion transport-related gene paralogs is consistent with the massive expansion of ion transport-related gene families in *E. affinis* complex genomes, relative to other arthropod and crustacean taxa (36,39,40). For example, 7-8 paralogs of *NHA* have been found in the *E. affinis* complex genome (36,39), whereas the genomes of other arthropods typically contain only 1-2 paralogs (25,44–47). In addition, *E. affinis* complex genomes contain 8 paralogs of *NKA-a*, 6 paralogs of *NKCC*, and 13 paralogs of *CA* (21,39,40,43).

Together, these previous results (previous paragraph) motivate further experiments to confirm, and more precisely quantify, the evolutionary and acclimatory expression changes for major ion transport-related genes. We were particularly interested in ion transport-related genes that showed allele frequency shifts (36,37) or evolutionary shifts in gene expression between saline and freshwater populations (29). In particular, 19 ion transport-related genes had previously shown salinity-induced evolutionary and/or acclimatory shifts in expression based on Illumina RNA-seq in *E. carolleeae* (29). However, targeted qPCR provides greater precision, better technical reproducibility, and improved ability to resolve gene expression differences among individual paralogs within expanded gene families (48–50).

Thus, this study employed quantitative real-time PCR (qPCR) to investigate evolutionary and acclimatory shifts in expression for these 19 ion transport-related genes in *E. carolleeae*. The specific aims of this study were to: **(1)** determine evolutionary shifts in ion transport-related gene expression and evolution of plasticity between saline and freshwater populations of *E. carolleeae* (34,51,52), **(2)** examine acclimatory changes in gene expression for these genes across salinities (0, 5, and 15 ppt) for each population, **(3)** compare the relative expression among the 19 ion transport-related genes (and their gene families) at two salinities (0 and 15 ppt) for each population, and **(4)** determine whether laboratory natural selection over six generations of salinity change recapitulates the evolutionary and plastic shifts observed between wild saline and freshwater populations.

To achieve the first and second goals, we performed a common garden reaction norm experiment, to determine genetically-based, heritable differences in gene expression between saline and freshwater populations at treatment salinities of 0, 5, and 15 ppt and acclimatory shifts across these salinities for each population. In this experiment, the saline ancestral and freshwater invading populations were first reared under “common garden” conditions (5 ppt) to remove acclimatory effects from their native salinities before being transferred to their common-garden reaction norm treatment salinities (0, 5, and 15 ppt). Resulting expression differences between the populations at each salinity (effect of *Population*) would represent heritable, genetically-based differences, while gene expression differences across treatment salinities (effect of *Salinity*) would represent plastic acclimatory responses for each population. A significant *Population* x *Salinity* interaction would indicate the evolution of plasticity between the saline and freshwater populations. To accomplish the third goal, we compared the relative expression of each ion transport-related gene family (and their paralogs) to one another for each population (saline and freshwater) and treatment salinity (0 or 15 ppt) (e.g., saline population reared at 0 ppt). To accomplish the fourth goal, we conducted a laboratory natural selection experiment to determine the evolutionary shifts in gene expression in response to salinity change over multiple generations. Specifically, we subjected an ancestral saline population from the St. Lawrence estuary to salinity decline (from 15 to 0 ppt) and a freshwater invading population from Lake Michigan to salinity increase (from 0 to 15 ppt) for six generations each.

Based on our previous genome-wide study (see above) (29), as well as our other ion transporter studies (21,23,30,31), we generated several hypotheses on the acclimatory and evolutionary responses of key ion transporter genes to salinity change. Specifically, we predicted that **(1)** certain ion transport-related genes will show evolutionary shifts in expression between ancestral saline and invading freshwater populations, such as paralogs of *NHA (NHA-5* and *NHA-7), NKA (NKA-α-1* and *NKA-α-2), NKCC (NKCC-2 and NKCC-3),* and *CA* (*CA-9*, *CA-12*, and *CA-11*) and **(2)** the paralogs *NHA-5, NHA-7,* and *NKCC-2* will show acclimatory shifts in response to salinity for both populations. In addition, while the prior genome-wide gene expression study did not examine the evolution of plasticity between the saline and freshwater populations (29), we predicted that **(3)** certain ion transport-related genes will exhibit the evolution of plasticity, with the freshwater population showing reduced plasticity (canalization) relative to the saline populations (53–55). In addition, based on our prior *in situ* expression studies (31), we predicted that **(4)** the level of expression among paralogs within gene families will differ for a given population at a given salinity, supporting the hypothesis of functional divergence among the paralogs. Regarding the laboratory natural selection experiment, we hypothesized that **(5)** the subset of genes showing the strongest differences in expression between populations in the common garden experiment will also be the most responsive to six generations of salinity induced laboratory natural selection. Furthermore, **(6)** we predicted that the genes showing evolutionary shifts in gene expression during laboratory selection will shift in the same direction as that observed between the wild saline and freshwater populations.

This study represents a detailed investigation of evolutionary and acclimatory shifts in ion transporter gene expression following rapid evolutionary transitions between salinities. The experiments performed here are notable in examining evolutionary shifts between saline and freshwater wild populations, the evolution of plasticity between these populations, as well as evolutionary and acclimatory responses following laboratory natural selection. Studies that examine the differential roles of individual ion transporter paralogs in contributing to salinity acclimation and adaptation have been extremely rare. These findings reveal not only how *E. carolleeae* repeatedly adapts to freshwater habitats, but also provide a framework for predicting species resilience under accelerating climate-driven salinity change (22).

## RESULTS

### 1. Evolutionary and acclimatory shifts in ion transporter gene expression under common garden conditions

Using a common garden experimental design (Fig 2A), analysis of gene expression patterns of 19 ion transport-related genes revealed both evolutionary shifts between the saline and freshwater populations and acclimatory responses across salinities (0, 5, and 15 ppt) for both populations, as well as the evolution of plasticity between the populations (Figs 3-5; Tables 1 and 2). Genes that showed the evolution of increased expression in the freshwater population relative to the ancestral saline population included *NKA-α-1* and *NKCC-3* (Fig 4). In contrast, *NHA-5* and *NHA-7* showed the evolution of increased expression in the saline population at 0 ppt (Figs 3 and 4; Table 1). Genes showing significant acclimatory responses across salinities for both populations included *NHA-5*, *NHA-7,* and *NKCC-2,* which showed significantly elevated expression at 0 ppt, and *CA-9*, which showed significantly higher expression at 15 ppt (Figs 3 and 4; Table 1). Genes showing the evolution of plasticity between saline and freshwater populations, based on significant *Population* x *Salinity* interaction effects, included *NKA-α-1, NHA-5, NHA-7, NKCC-2*, and *CA-9* (Figs 4 and S4; S3 Table).

**Fig 2.**
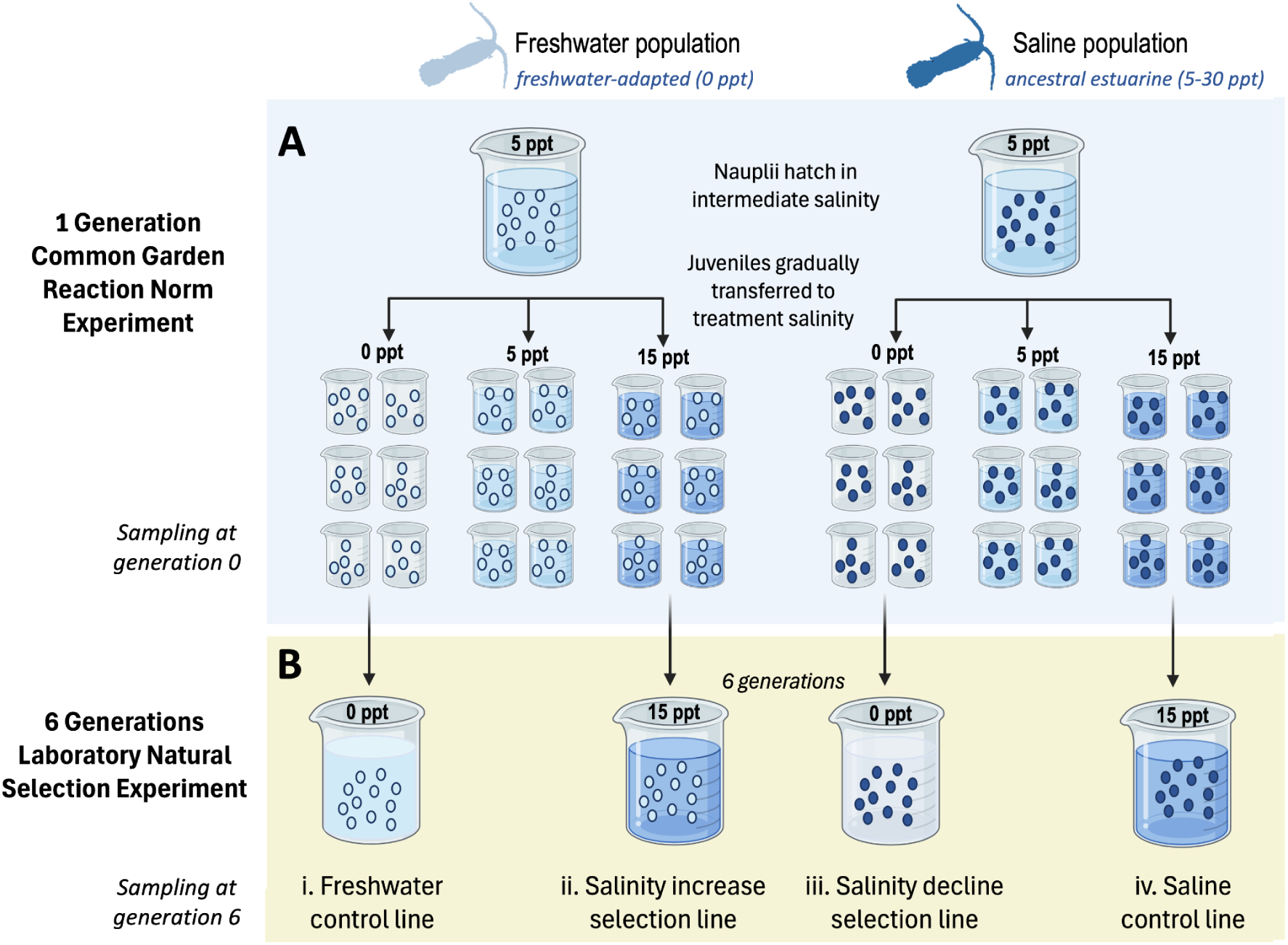
Design of laboratory common garden reaction norm and natural selection experiments using wild saline and freshwater populations of *Eurytemora carolleeae*. The ancestral saline population is from the St. Lawrence estuary, whereas the derived freshwater population is from Lake Michigan (Fig 1A). (**A**) Common garden reaction norm experiment to determine heritable differences in ion transporter gene expression between the saline and freshwater populations and acclimatory responses across salinities (0, 5, 15 ppt) after one generation of rearing at each treatment salinity. The figure shows only one block out of two replicate blocks of the experiment. A single block contained six biological replicates (replicate beakers containing ∼50 copepods) for each salinity treatment for each population, for a total of twelve biological replicates across both blocks per salinity treatment per population. (**B**) Laboratory natural selection experiments, imposing selection for salinity increase in the freshwater population and salinity decline in the saline population. The freshwater population was subjected to salinity increase, from the control salinity of 5 ppt to 15 ppt, within one generation (during the common garden experiment) and then remained at 15 ppt for six generations (Line ii). The saline population was subjected to salinity decline, from 5 to 0 ppt, within one generation (during the common garden experiment) and then remained at 0 ppt for six generations ( Line iii). Both selection lines were compared to control lines for each population maintained at their native salinities for the duration of the experiment, namely 0 ppt for the freshwater population (Line i) and 15 ppt for the saline population (Line iv). One beaker was maintained for each experimental line. From each beaker 4-6 biological replicates were sampled, with each replicate consisting of a pool of 50 animals. The number of replicates sampled per beaker was limited by the number of animals surviving in each salinity (Line i, N=4 replicates; Line ii, N=6; Line iii, N=4; Line iv, N=6).

**Table 1.** Ion transport-related genes exhibiting **(A–E)** evolutionary shifts in expression between saline (SW) and freshwater (FW) populations, as well as **(F–J)** acclimatory shifts in expression across common-garden salinity treatments (0 and 15 ppt) within each population. Mean expression values are based on N =12 biological replicates per population and salinity treatment (Fig 2A). Values shown for evolutionary comparisons are **(A)** treatment salinity in which populations were compared, **(B)** Mean ± SE expression for the saline population, **(C)** Mean ± SE expression for the freshwater population, **(D)** Log_2_ fold change in expression (the ratio) between the freshwater *vs.* saline population (Log_2_(FC): FW/SW). Positive Log_2_fold change values indicate higher expression in the freshwater population relative to the saline population. **(E)** *P*-values from Tukey HSD *post hoc* tests between the saline and freshwater population values, following a two-way ANOVA (S1 and S3 Tables). Values shown for the acclimatory comparisons are shown for **(F)** population for which treatment salinities were compared, **(G)** Mean ± SE expression at 0 ppt, **(H)** Mean ± SE expression at 15 ppt, **(I)** Log_2_ fold change in expression (the ratio) between the 0 *vs.* 15 ppt treatment salinities (Log_2_(FC): 0/15 ppt). Positive Log_2_ fold change values indicate higher expression in the 0 ppt treatment relative to the 15 ppt salinity. **(J)** *P*-values from Tukey HSD post hoc tests between the 0 and 15 ppt values following a two-way ANOVA (S2 and S3 Tables). Significance levels are indicated as \**P* < 0.05, \*\**P* < 0.01, \*\*\**P* < 0.001, \*\*\*\**P* < 0.0001. The full list of genes with two-way ANOVA results are provided in S3 Table and corresponding Tukey HSD post hoc tests are provided in S1, and S2 Tables.

| **Gene Name** | **(A)<br>Salinity** | **(B)<br>SW pop mean ± SE** | **(C)<br>FW pop mean ± SE** | **(D)<br>Log<sub>2</sub>(FC):<br>FW pop/SW pop** | **(E)<br>Tukey HSD *P*-value** |
|---|---|---|---|---|---|
| *NKA-α-1* | 0 | 1.16 ± 0.277 | 18.98 ± 2.24 | 4.04 | 2.84 x 10<sup>-10</sup>**** |
|  | 15 | 0.833 ± 0.147 | 15.03 ± 1.90 | 4.17 | 3.51 x 10<sup>-9</sup>**** |
| *NKCC-3* | 0 | 0.875 ± 0.104 | 1.79 ± 0.105 | 1.03 | 0.0419* |
|  | 15 | 0.524 ± 0.138 | 1.30 ± 0.118 | 1.31 | 0.0530* |
| *NHA-5* | 0 | 6.61 ± 1.29 | 0.139 ± 0.079 | -5.57 | 0.00116*** |
|  | 15 | 0.639 ± 0.0710 | 0.180 ± 0.062 | -1.83 | 0.999 |
| **Gene Name** | **(F)<br>Population** | **(G)<br>0 ppt mean ± SE** | **(H)<br>15 ppt mean ± SE** | **(I)<br>Log<sub>2</sub>(FC):<br>0/15 ppt** | **(J)<br>Tukey HSD *P*-value** |
| *NKCC-2* | FW | 5.11 ± 0.435 | 1.88 ± 0.082 | 1.45 | 3.23 x 10<sup>−7</sup>**** |
|  | SW | 3.84 ± 0.348 | 0.204 ± 0.021 | 4.24 | 9.30 x 10<sup>-11</sup>**** |
| *NHA-7* | FW | 1.65 ± 0.107 | 0.803 ± 0.065 | 1.04 | 0.0833 |
|  | SW | 4.58 ± 0.334 | 0.616 ± 0.114 | 2.90 | 8.25 x 10<sup>−13</sup>**** |
| *CA-9* | FW | 0.452 ± 0.071 | 3.56 ± 0.431 | -2.98 | 6.95 x 10<sup>−5</sup>**** |
|  | SW | 0.108 ± 0.041 | 2.39 ± 0.523 | -4.47 | 0.00133*** |

**Table 2.**
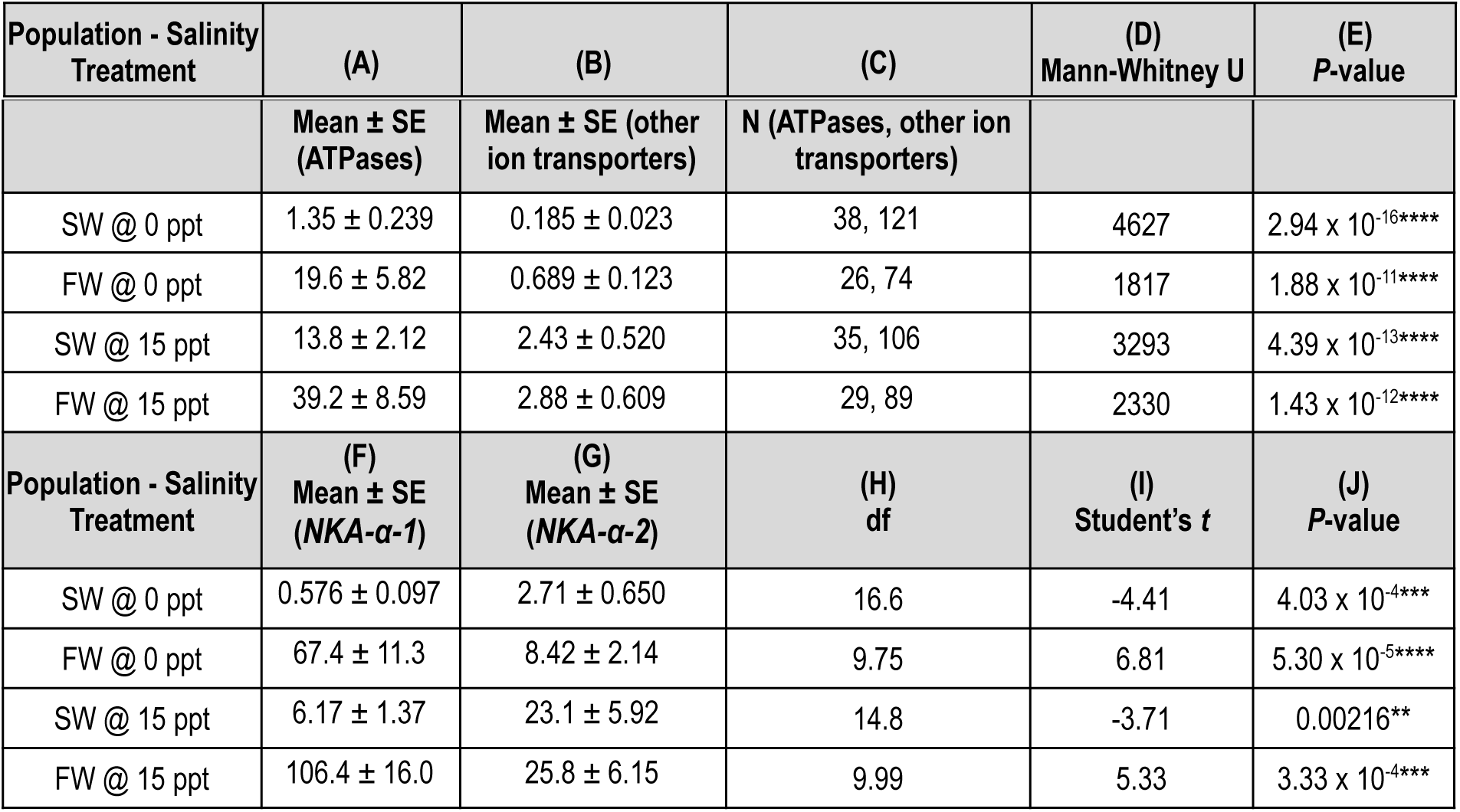
Comparison of gene expression between ATPases (*NKA-α-1*, *NKA-α-2*, *VHA-a* and *VHA-H*) *vs.* other ion transporter-related genes and between *NKA-α-1 vs*. *NKA-α-2*. Comparisons were made for each population (SW or FW) reared at two treatment salinities (0 or 15 ppt). A pooled Mann-Whitney-U test was used to test the differences in gene expression between four ATPases *vs.* 15 other ion transport-related genes, because the groups differed greatly in expression level and sample size. For each population (SW or FW) at each treatment salinity (0 or 15 ppt) **(A)** Mean ± SE of gene expression of four ATPase genes (*NKA-α-1*, *NKA-α-2*, *VHA-a* and *VHA-H*), **(B)** Mean ± SE of gene expression of the other 15 ion transport-related genes, **(C)** Sample size for each group (total number of replicate measurements for the ATPases, followed by the total number of replicates for all other ion transport-related genes, **(D)** test statistic for the Mann-Whitney test (U-value), and **(E)** *P*-value indicating the level of significant difference for each comparison, indicated by asterisks within the table (**** *P* < 0.0001). A Student’s t-test was used to test the differences in gene expression between the NKA paralogs ( *NKA-α-1* vs. *NKA-α-2*) for each population (SW or FW) reared at two treatment salinities (0 or 15 ppt). For each population (SW or FW) at each treatment salinity (0 or 15 ppt) **(F)** Mean ± SE of gene expression of *NKA-α-1*, **(G)** Mean ± SE of gene expression of *NKA-α-2*, **(H)** degrees of freedom (df) for the comparison **(I)** test statistic for the Student’s t-test (t), **(J)**, and the associated *P*-value and level of significance, indicated by asterisks within the graph (\*\**P* < 0.01, \*\*\**P* < 0.001, \*\*\*\**P* < 0.0001).

Of the 19 osmo/ionoregulatory genes examined, we identified single nucleotide polymorphisms (SNPs) between the saline and freshwater populations in the PCR primer sequences of three genes: *NHA-1*, *NHA-3*, and *NHA-6* (Table 4). These SNPs could potentially cause population-specific primer annealing biases. Because standard curves were generated using pooled cDNA from both populations, we could not verify whether primer efficiency for these three genes was equivalent for the saline and freshwater populations. Therefore, these three genes were excluded from the between-population expression comparisons but were retained for within-population analyses of acclimatory responses across salinities. Importantly, no significant block effects for replicate experiments (Blocks 1 and 2) were detected for any of the genes, indicating that experimental block differences did not influence gene expression patterns.

For all other genes examined, for which SNPs between populations were not present in the PCR primer sequences, we compared gene expression patterns between populations (saline *vs.* fresh) at different salinities (0, 5, and 15 ppt). For the expression levels of these genes, we performed a Principal Component Analysis (PCA) to identify general patterns of gene expression between populations and across salinities. The first principal component (PC1) accounted for 38.1% of the variance, while PC2 and PC3 explained 26.8% and 16.9% of the variance, respectively (Fig 3).

**Fig 3.**
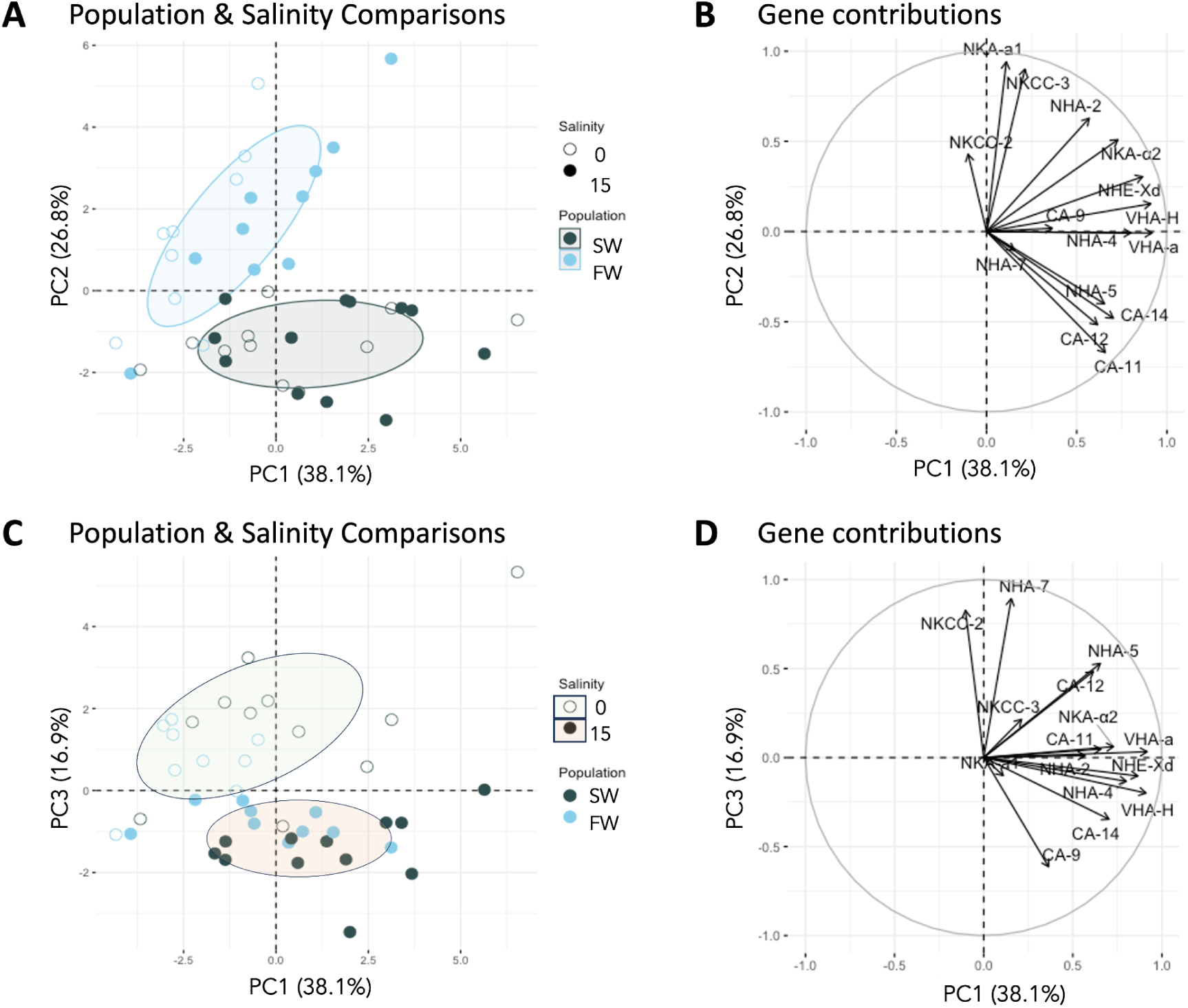
Principal Component Analysis (PCA) of relative expression for 15 ion transport-related genes of *E. carolleeae* populations from the common garden experiment of saline and freshwater populations reared at three salinities (see Fig 2A). “Relative expression” here refers to the amount of ion transporter (and CA) mRNA expression relative to the mean expression of housekeeping genes (*EF-1α* and *GDPH*) (Equation 1 in Methods). (**A**) Plot of the PC1 and PC2 coordinate plane showing the PC scores for the gene expression profiles from saline (SW, black dots) and freshwater (FW, blue dots) populations reared under common garden treatment salinities of 0 ppt (open dots) and 15 ppt (closed dots). Each data point is one biological replicate (a separate beaker each containing a pool of 50 animals) (N = 12 biological replicates per population and salinity treatment, Fig 2A). Ellipses represent 50% confidence intervals, chosen to illustrate the core spread of each group while minimizing overlap and emphasizing differences in group tightness and separation. (**B**) Circle plot showing the contributions of each gene’s expression to the PC1 and PC2 components. Longer arrows indicate larger contribution and direction of arrows reflects the correlation between a gene’s expression pattern and the PCA axes. *Population* had a significant effect on gene expression patterns (PCA scores) (two-way MANOVA; Pillai’s Trace = 0.671, *F*(2,37) = 37.5, *P* = 1.16 x 10^-9^), mainly along PC2, indicating evolutionary shifts in ion transport-related gene expression between the populations. **(C)** PC1 and PC3 coordinate plane showing the PC scores for gene expression profiles from saline (SW, black dots) and freshwater (FW, blue dots) populations reared under common garden treatment salinities of 0 ppt (open dots) and 15 ppt (closed dots). Each data point is one biological replicate (a separate beaker each containing a pool of 50 animals) (N = 12 biological replicates per population and salinity treatment, Fig 2A). Ellipses represent 50% confidence intervals. **(D)** Circle plot showing the contributions of each gene’s expression to the PC1 and PC3 components. *Salinity* had a significant effect on gene expression patterns (PCA scores) (two-way MANOVA Pillai’s Trace = 0.649, *F*(2,37) = 34.2, *P* = 3.87 x 10^-9^), indicating acclimatory shifts in ion transport-related gene expression between salinities.

The PCA revealed clear differences in ion transport-related gene expression patterns between saline and freshwater populations, as shown in the PC1 and PC2 coordinate plane (Fig 3A) and across salinities, as displayed in the PC1 and PC3 coordinate plane (Fig 3C). For each PCA plot, a multivariate analysis of variance (Two-way MANOVA) was performed to determine the effects of the factors *Population* (freshwater *vs.* saline populations), *Salinity* (0 *vs.* 15 ppt), and *Population* x *Salinity* interaction on the response variable *PCAscores* of the data points (shown in Fig 3A and 3C). For this analysis, each data point on the PCA plots (Fig 3) represented an individual biological replicate for each population (saline and freshwater) at each salinity (0 and 15 ppt) from the common garden experiment (N = 12 per treatment and per population).

Our results indicated the presence of evolutionary shifts in gene expression between saline and freshwater populations (Fig 3A and B). In the PC1 and PC2 coordinate plane (Fig 3A), the saline and freshwater populations formed separate clusters. Consistent with this pattern, the factor *Population* had a significant effect on *PCAscores* of individual data points (two-way MANOVA; Pillai’s Trace = 0.671, *F*(2,37) = 37.5, *P* = 1.16 x 10^-9^), indicating evolutionary shifts between the populations. The effect of *Salinity* on *PCAscores* was marginally significant in this coordinate plane (two-way MANOVA; Pillai’s Trace = 0.208, *F*(2,37) = 4,85, *P* = 0.0134). *Population x Salinity* effects on *PCAscores* were not significant (two-way MANOVA; Pillai’s Trace = 0.012, *F*(2,37) = 0.258, *P* = 0.774).

As mentioned above, ion transport-related gene expression between saline and freshwater populations separated clearly along the PC2 axis (Fig 3A, blue *vs.* black circles), reflecting evolutionary shifts between them. Gene clustering along the PC2 axis (Fig 3B) included genes potentially contributing to adaptive divergence in ion transport function between saline and freshwater populations (21,22,29). Positive values along PC2 for *NKA-α-1, NKCC-3* and to a lesser extent *NKCC-2* (Fig 3B) indicated the evolution of increased gene expression in the freshwater population relative to the ancestral saline population (Figs 4, S3, and S4; Table 1). This evolution of increased expression in the freshwater population was significant for *NKCC-3* and *NKA-α-1* across all salinities and for *NKCC-2* at 15 ppt (Fig 4; Tables 1 and S1). On the other hand, negative values along PC2 for the genes *CA-11, CA-12, CA-14, NHA-5,* and to a much lesser extent *NHA-7* (Fig 3B) indicated the evolution of increased expression in saline populations (Figs 4, S3 and S4; Table 1). This trend of evolution of higher expression in the saline population was significant for *CA-12* at 0 and 5 ppt, for *CA-11* at 0 and 15 ppt, and for *NHA-5* and *NHA-7* at 0 ppt (Figs 4 and S4; Tables 1 and S1).

**Fig 4.**
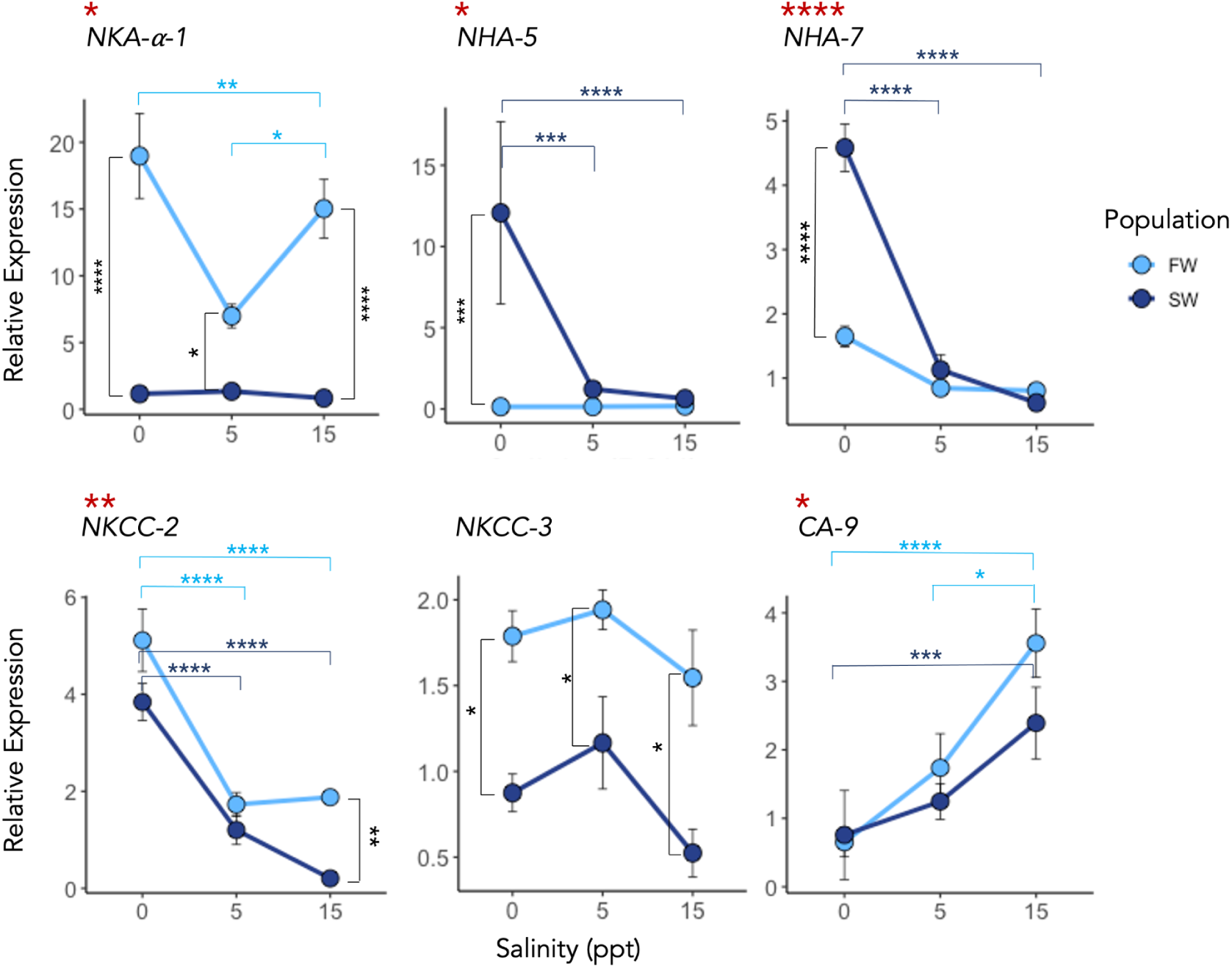
Evolutionary shifts in expression of six ion transport-related genes between saline (SW) and freshwater (FW) populations and acclimatory shifts across the salinities (0, 5, and 15 ppt) under common garden conditions (see Fig 2A). “Relative expression” here refers to the amount of ion transporter (and CA) mRNA expression relative to the mean expression of housekeeping genes (*EF-1α* and *GDPH*) (Equation 1 in Methods). Data are represented as Mean ± SE across 12 biological replicates (Fig 2A). Statistical analyses were conducted using a two-way ANOVA. Red **\*** near the gene names indicates a significant interaction effect between *Population* x *Salinity* (i.e. evolution of plasticity) and significant results from TukeyHSD *post hoc* analysis are indicated by asterisks within each graph (\**P* < 0.05, \*\**P* < 0.01, \*\*\**P* < 0.001, \*\*\*\**P* < 0.0001). Expression patterns for all genes are shown in S3, S4, and S5 Figs. Statistical results are reported in Tables 1, S1, S2, and S3.

Also in the PC1 and PC2 coordinate plane (Fig 3A), the PC1 axis was associated with a subtle acclimatory response in expression between salinity treatments (Fig 3A, open *vs.* closed circles), consistent with the marginally significant effect of *Salinity* on *PCAscores* in this coordinate plane (see above). This acclimatory response was mainly attributed to changes in expression of *VHA-H, VHA-α, NHA-4*, *NHE-Xd* that clustered together along the PC1 axis (Fig 3B). These genes, as well as *CA-9,* showed positive values along the PC1 axis (Figs 3B, 3D, and S3). In both populations their expression tended to exhibit lower expression at lower salinity and higher expression at higher salinity, however, only *CA-9* showed a significant effect (Figs 4 and S4).

In the PC1 and PC3 coordinate plane (Fig 3C), a substantial acclimatory response was evident in ion transport-related gene expression, with clear separation of data points between treatment salinities 0 and 15 ppt (Fig 3C, open *vs.* closed circles), particularly along the PC3 axis. This acclimatory response was evident from the highly significant effect of *Salinity* on *PCAscores* of the data points (two-way MANOVA; Pillai’s Trace = 0.649, *F*(2,37) = 37.5, *P* = 3.87 x 10^-9^). In contrast, the effect of *Population* on *PCAscores* was not significant on this coordinate plane (two-way MANOVA; Pillai’s Trace = 0.111, *F*(2,37) = 2.31, *P* = 0.113). *Population x Salinity* interaction effects on *PCAscores* were also not significant (two-way MANOVA; Pillai’s Trace = 0.132, *F*(2,37) = 2.82, *P* = 0.073). Positive values along the PC3 axis for *NHA-7* and *NKCC-2* (Fig 3D) reflected the increase in gene expression at low salinity (0 ppt) relative to higher treatment salinities (15 ppt) for both the saline and freshwater populations (Figs 4 and S4; Table 3). In contrast, negative values along the PC3 axis (Fig 3D) indicated higher gene expression in the 15 ppt relative to 0 ppt treatment, such as for *CA-9* and *CA-14,* which both clustered together in the lower right quadrant of Figure 4D (Figs 4 and S4; Tables 3 and S2).

**Table 3.**
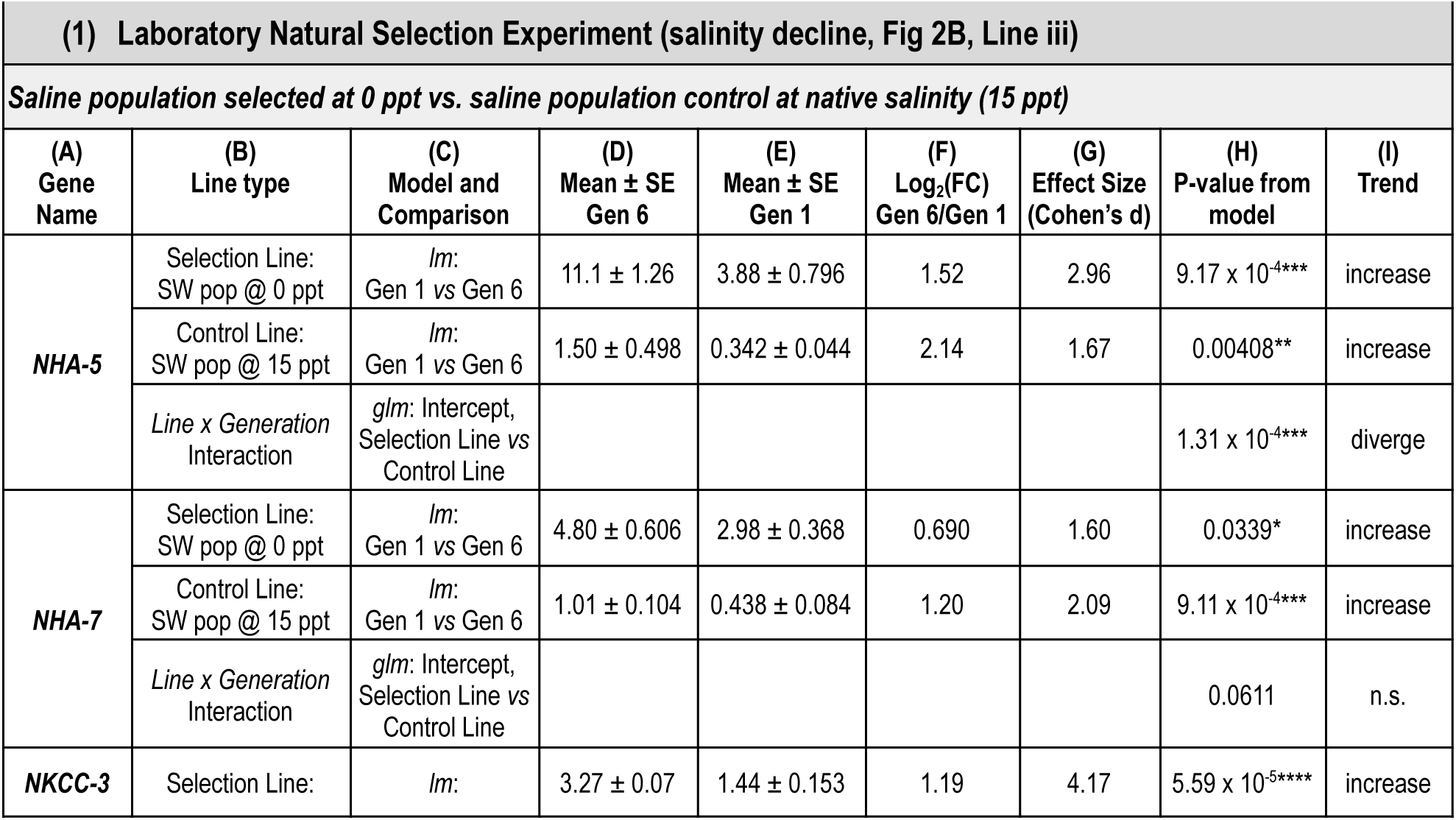
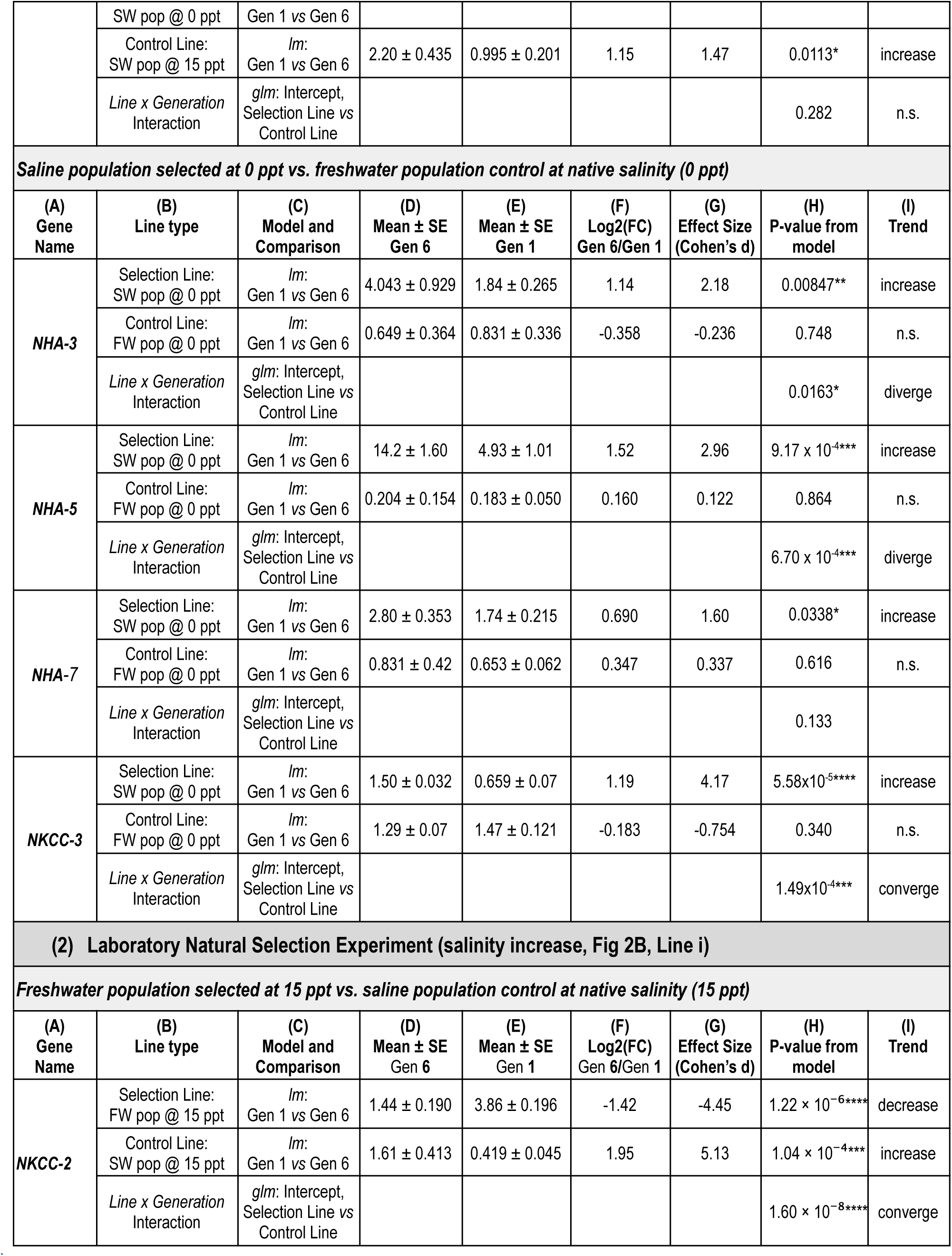
Comparison of gene expression trajectories between laboratory natural selection lines *versus* control lines (see Fig 2B). The table includes two main experimental designs: **(1)** a saline population of *Eurytemora carolleeae* undergoing salinity decline (selected at 0 ppt for six generations, Fig 2B, Line iii), relative to two control lines. The control lines consisted of a saline population maintained at native salinity (15 ppt, Line iv) and a freshwater population maintained at native salinity (0 ppt, Line i); and **(2)** a freshwater population undergoing salinity increase (selected at 15 ppt for six generations, Fig 2B, Line i), relative to the same two control lines as in (1). Within each experimental design, columns show **(A)** the gene name, **(B)** the type of experimental line (selection line or control line) and its corresponding population and salinity, **(C)** the statistical model and comparison, using either a linear model (*lm*) for the within-line shift across generations (Generation 1 *vs.* Generation 6) or a generalized linear model (glm) for the between-line comparison (Selection line *vs.* Control line). Values shown are the Mean ± SE expression values for the number of animals surviving at each salinity shown in Fig 2B for **(D)** Generation 6 and **(E)** Generation 1, **(F)** Log_2_ fold change (Log2(FC): Gen 6/Gen 1), and **(G)** effect size (Cohen’s *d*), which quantifies the direction, magnitude, and standardized strength of expression shifts. **(H)** *P*-values obtained from linear models (lm) for within-line comparisons and generalized linear models (glm) for between-line comparisons. Significance levels are indicated as * *P* < 0.05, \*\**P* < 0.01, \*\*\**P* < 0.001, \*\*\*\**P* < 0.0001. The **(I)** Trend reflects either the direction of expression shift (increase or decrease) or whether the two trajectories are diverging or converging (see Fig 6).

| **(1) Laboratory Natural Selection Experiment (salinity decline, Fig 2B, Line iii)** |  |  |  |  |  |  |  |  |
|---|---|---|---|---|---|---|---|---|
| ***Saline population selected at 0 ppt vs. saline population control at native salinity (15 ppt)*** |  |  |  |  |  |  |  |  |
| **(A)<br>Gene<br>Name** | **(B)<br>Line type** | **(C)<br>Model and<br>Comparison** | **(D)<br>Mean ± SE<br>Gen 6** | **(E)<br>Mean ± SE<br>Gen 1** | **(F)<br>Log<sub>2</sub>(FC)<br>Gen 6/Gen 1** | **(G)<br>Effect Size<br>(Cohen's d)** | **(H)<br>P-value from<br>model** | **(I)<br>Trend** |
| ***NHA-5*** | Selection Line:<br>SW pop @ 0 ppt | *lm*:<br>Gen 1 *vs* Gen 6 | 11.1 ± 1.26 | 3.88 ± 0.796 | 1.52 | 2.96 | 9.17 x 10<sup>-4</sup>*** | increase |
|  | Control Line:<br>SW pop @ 15 ppt | *lm*:<br>Gen 1 *vs* Gen 6 | 1.50 ± 0.498 | 0.342 ± 0.044 | 2.14 | 1.67 | 0.00408** | increase |
|  | *Line x Generation* Interaction | *glm*: Intercept, Selection Line *vs* Control Line |  |  |  |  | 1.31 x 10<sup>-4</sup>*** | diverge |
| ***NHA-7*** | Selection Line:<br>SW pop @ 0 ppt | *lm*:<br>Gen 1 *vs* Gen 6 | 4.80 ± 0.606 | 2.98 ± 0.368 | 0.690 | 1.60 | 0.0339* | increase |
|  | Control Line:<br>SW pop @ 15 ppt | *lm*:<br>Gen 1 *vs* Gen 6 | 1.01 ± 0.104 | 0.438 ± 0.084 | 1.20 | 2.09 | 9.11 x 10<sup>-4</sup>*** | increase |
|  | *Line x Generation* Interaction | *glm*: Intercept, Selection Line *vs* Control Line |  |  |  |  | 0.0611 | n.s. |
| ***NKCC-3*** | Selection Line:<br>SW pop @ 0 ppt | *lm*:<br>Gen 1 *vs* Gen 6 | 3.27 ± 0.07 | 1.44 ± 0.153 | 1.19 | 4.17 | 5.59 x 10<sup>-5</sup>**** | increase |
|  | Control Line:<br>SW pop @ 15 ppt | *lm*:<br>Gen 1 *vs* Gen 6 | 2.20 ± 0.435 | 0.995 ± 0.201 | 1.15 | 1.47 | 0.0113* | increase |
|  | *Line x Generation* Interaction | *glm*: Intercept, Selection Line *vs* Control Line |  |  |  |  | 0.282 | n.s. |
| ***Saline population selected at 0 ppt vs. freshwater population control at native salinity (0 ppt)*** |  |  |  |  |  |  |  |  |
| **(A)<br>Gene<br>Name** | **(B)<br>Line type** | **(C)<br>Model and<br>Comparison** | **(D)<br>Mean ± SE<br>Gen 6** | **(E)<br>Mean ± SE<br>Gen 1** | **(F)<br>Log2(FC)<br>Gen 6/Gen 1** | **(G)<br>Effect Size<br>(Cohen's d)** | **(H)<br>P-value from<br>model** | **(I)<br>Trend** |
| ***NHA-3*** | Selection Line:<br>SW pop @ 0 ppt | *lm*:<br>Gen 1 *vs* Gen 6 | 4.043 ± 0.929 | 1.84 ± 0.265 | 1.14 | 2.18 | 0.00847** | increase |
|  | Control Line:<br>FW pop @ 0 ppt | *lm*:<br>Gen 1 *vs* Gen 6 | 0.649 ± 0.364 | 0.831 ± 0.336 | -0.358 | -0.236 | 0.748 | n.s. |
|  | *Line x Generation* Interaction | *glm*: Intercept, Selection Line *vs* Control Line |  |  |  |  | 0.0163* | diverge |
| ***NHA-5*** | Selection Line:<br>SW pop @ 0 ppt | *lm*:<br>Gen 1 *vs* Gen 6 | 14.2 ± 1.60 | 4.93 ± 1.01 | 1.52 | 2.96 | 9.17 x 10<sup>-4</sup>*** | increase |
|  | Control Line:<br>FW pop @ 0 ppt | *lm*:<br>Gen 1 *vs* Gen 6 | 0.204 ± 0.154 | 0.183 ± 0.050 | 0.160 | 0.122 | 0.864 | n.s. |
|  | *Line x Generation* Interaction | *glm*: Intercept, Selection Line *vs* Control Line |  |  |  |  | 6.70 x 10<sup>-4</sup>*** | diverge |
| ***NHA-7*** | Selection Line:<br>SW pop @ 0 ppt | *lm*:<br>Gen 1 *vs* Gen 6 | 2.80 ± 0.353 | 1.74 ± 0.215 | 0.690 | 1.60 | 0.0338* | increase |
|  | Control Line:<br>FW pop @ 0 ppt | *lm*:<br>Gen 1 *vs* Gen 6 | 0.831 ± 0.42 | 0.653 ± 0.062 | 0.347 | 0.337 | 0.616 | n.s. |
|  | *Line x Generation* Interaction | *glm*: Intercept, Selection Line *vs* Control Line |  |  |  |  | 0.133 |  |
| ***NKCC-3*** | Selection Line:<br>SW pop @ 0 ppt | *lm*:<br>Gen 1 *vs* Gen 6 | 1.50 ± 0.032 | 0.659 ± 0.07 | 1.19 | 4.17 | 5.58x10<sup>-5</sup>**** | increase |
|  | Control Line:<br>FW pop @ 0 ppt | *lm*:<br>Gen 1 *vs* Gen 6 | 1.29 ± 0.07 | 1.47 ± 0.121 | -0.183 | -0.754 | 0.340 | n.s. |
|  | *Line x Generation* Interaction | *glm*: Intercept, Selection Line *vs* Control Line |  |  |  |  | 1.49x10<sup>-4</sup>*** | converge |
| **(2) Laboratory Natural Selection Experiment (salinity increase, Fig 2B, Line i)** |  |  |  |  |  |  |  |  |
| ***Freshwater population selected at 15 ppt vs. saline population control at native salinity (15 ppt)*** |  |  |  |  |  |  |  |  |
| **(A)<br>Gene<br>Name** | **(B)<br>Line type** | **(C)<br>Model and<br>Comparison** | **(D)<br>Mean ± SE**<br>Gen **6** | **(E)<br>Mean ± SE**<br>Gen **1** | **(F)<br>Log2(FC)**<br>Gen **6**/Gen **1** | **(G)<br>Effect Size<br>(Cohen's d)** | **(H)<br>P-value from<br>model** | **(I)<br>Trend** |
| ***NKCC-2*** | Selection Line:<br>FW pop @ 15 ppt | *lm*:<br>Gen 1 *vs* Gen 6 | 1.44 ± 0.190 | 3.86 ± 0.196 | -1.42 | -4.45 | 1.22 × 10<sup>−6</sup>**** | decrease |
|  | Control Line:<br>SW pop @ 15 ppt | *lm*:<br>Gen 1 *vs* Gen 6 | 1.61 ± 0.413 | 0.419 ± 0.045 | 1.95 | 5.13 | 1.04 × 10<sup>−4</sup>*** | increase |
|  | *Line x Generation* Interaction | *glm*: Intercept, Selection Line *vs* Control Line |  |  |  |  | 1.60 × 10<sup>−8</sup>**** | converge |

In addition to evolutionary shifts between saline and freshwater populations (Figs 3, 4, and S4; Tables 1 and S1), we also observed the evolution of plasticity between the populations (Figs 4 and S4; S3 Table). The evolution of plasticity between the saline and freshwater populations was evident from the significant change in slope of the acclimatory (plastic) response across salinities (shift in reaction norms) between the populations (graphs with red stars in Figs 4, and S4). This evolution in slope between the populations was indicated by the significant *Population x Salinity* interaction term in the two-way ANOVA (S3 Table). The ion transport-related genes that displayed the evolution of plasticity between the populations included *NHA-7, NHA-5, NKA-α-1, NKCC-2*, and *CA-9* (Fig 4). In the cases of *NHA-7* and *NHA-5*, these genes exhibited greater plasticity in the ancestral saline populations followed by the evolution of canalization in the freshwater population (Figs 4, 6, and S4; Table 1).

### 2. Variation in expression patterns among the ion transport-related genes

To determine variation in expression patterns among the ion transport-related genes, we compared the expression levels of 19 genes within each population at each salinity treatment (Fig 5). Specifically, we compared the expression levels of individual ion transport-related genes and functional gene groups within each population (i.e., saline and freshwater populations) at each treatment salinity (i.e., 0 and 15 ppt) (Fig 2A and 5). Across populations and salinities, the ion pumps (ATPases: *NKA* paralogs and *VHA* subunits) tended to show the highest expression levels (Fig 5A–D; light orange shading). When analyzed as groups, these ATPases showed significantly higher gene expression compared to all other ion transport-related genes (Table 2).

**Fig 5.**
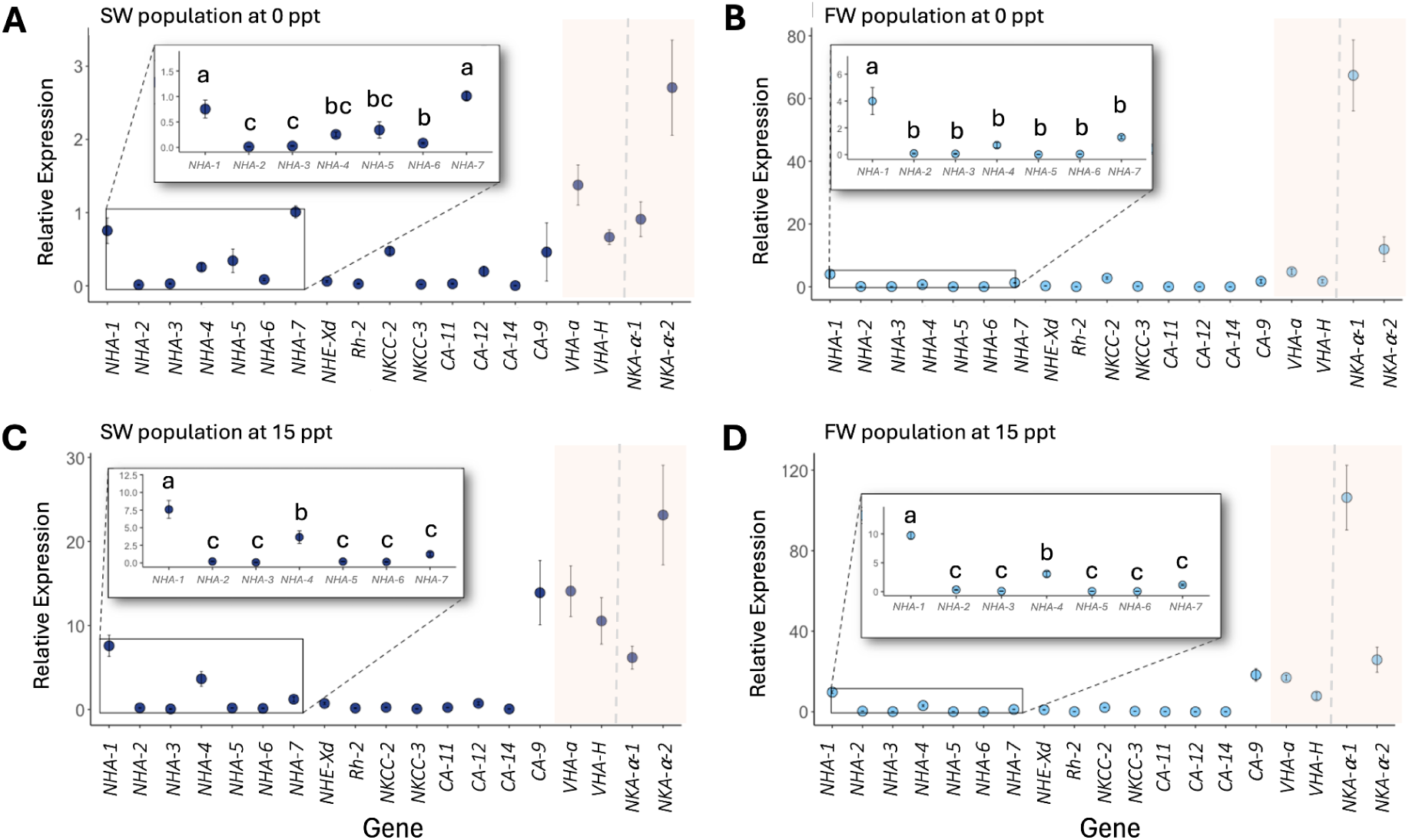
Relative expression of ionoregulatory genes in *E. carolleeae* from ancestral saline (SW) and invading freshwater (FW) populations, following one generation of rearing under treatment salinities of 0 or 15 ppt during a common garden reaction norm experiment (see Fig 2A). “Relative expression” here refers to the amount of ion transporter (and CA) transcript level relative to the mean expression of housekeeping genes ( *EF-1α* and *GDPH*) (Equation 1 in Methods). Data points are shown as Mean ± SE (Standard Error). Insets magnify the relative expression of the *NHA* (Na^+^/H^+^ antiporter) paralogs. Pairwise comparisons among the NHA paralogs were conducted using Student’s t-tests with a Bonferroni correction; lower case letters represent statistical relationships among the *NHA* genes, where different letters indicate significant difference (statistical results shown in S4 Table). The light orange shading highlights the ATPases (VHA and NKA), which were compared to all other genes using a pooled Mann-Whitney test (shown in Table 2). The vertical dotted grey lines demarcate the *VHA* subunits from the two *NKA* paralogs. The *NKA* paralogs were compared to each other using a Student’s t-test (statistical results shown in Table 2).

To further examine variation within the *NKA* gene family, we then compared the expression patterns between the two *NKA* paralogs across populations and salinities. Within the saline population, the expression level of *NKA-α-2* displayed the highest expression among all the genes at both 0 and 15 ppt (Figs 5A and 5C; right of the dotted grey line). Between the two NKA paralogs, *NKA-α-2* showed ∼5x greater expression than *NKA-α-1* at both 0 and 15 ppt (Table 2). In contrast, *NKA-α-1* was the most highly expressed of all the genes in the freshwater populations under both salinity treatments (Figs 5B and 5D; right of the dotted grey line), with *NKA-α-1* showing 8x greater expression than *NKA-α-2* at 0 ppt and 4x greater expression than *NKA-α-2* at 15 ppt (Table 2).

Among the NHA paralogs, expression levels differed across population and salinity treatments (Fig 5A-D). *NHA-1* had the highest expression among the NHA paralogs, except in the saline population reared at 0 ppt (Fig 5A-D; S4 Table). Specifically, in the saline population at 0 ppt, both *NHA-1* and *NHA-7* showed significantly higher expression relative to the other NHA paralogs (Fig 5A). In addition, in the freshwater population at 0 ppt, *NHA-7* also had a trend toward greater expression, though not significant, than the NHA paralogs 2-6 (Fig 5A and 5B; S4 Table). Notably, the higher expression of *NHA-7* at 0 ppt (Fig 4) and higher expression relative to other NHA paralogs in the saline population at 0 ppt (Figs 5A and 5B; S4 Table) suggests its importance under low salinity conditions. In contrast, under 15 ppt conditions *NHA-4* showed the next highest expression after *NHA-1*, with significantly higher levels than the other NHA paralogs in both saline and freshwater populations (Figs 7C and 7D; S4 Table).

### 3. Evolutionary shifts in gene expression following laboratory selection in response to salinity change

Following the common garden experiment, we conducted a laboratory natural selection experiment to examine the evolutionary shifts in ion transporter (and carbonic anhydrase) gene expression in saline and freshwater populations of *E. carolleeae* following salinity change (Fig 2B). We established two selection lines, where the saline population experienced salinity decline and the freshwater population underwent salinity increase. For the salinity decline selection line, we gradually transferred the saline population (from St. Lawrence saltmarsh) from 15 ppt to 5 ppt to 0 ppt, following the step-wise salinity decline regime of the common garden experiment (5 → 2.5 → 1 → 0 ppt, with transfer to a new salinity every two days), within one generation, and then maintained the selection line at 0 ppt (lake water) for six generations (Fig 2B, line iii). For the salinity increase selection line, we gradually transferred the freshwater population (from Lake Michigan) from 0 ppt (lake water) to 5 ppt to 15 ppt, following the step-wise salinity increase regime of the common garden experiment (5 → 8 → 10 → 15 ppt, one transfer every two days), within one generation, and then maintained the line at 15 ppt for six generations (Fig 2B, line ii). We compared changes in mRNA levels between these selection lines and two control lines, which were the saline and freshwater populations maintained at their native salinities (15 and 0 ppt, respectively). For instance, we compared the saline population selected at 0 ppt to both control lines to assess whether gene expression patterns diverged from the ancestral saline population and converged toward expression patterns of the derived freshwater population. Similarly, we compared the freshwater population selected at 15 ppt to these same control lines to determine whether expression shifted away from the freshwater population and converged toward expression patterns of the ancestral saline population.

In the salinity decline experiment, the saline selection line at 0 ppt (Fig 2B, Line iii) was compared to the saline control line maintained at its native 15 ppt salinity (Fig 2B, Line iv). First, within-line comparisons between Generation 1 and Generation 6 in the saline selection line (Line iii) showed that gene expression of *NHA-5*, *NHA-7*, and *NKCC-3* increased significantly at Generation 6 at 0 ppt (approximately 3-fold, 1.6-fold, and 2-fold, respectively; Fig 6A, Table 3). In the 15 ppt control line (Line iv), gene expression trajectory of *NHA-5*, *NHA-7* and *NKCC-3* also increased significantly across generations (Fig 6A, Table 3). Of these comparisons, only *NHA-5*, showed a significant *Line* x *Salinity* interaction effect (Fig 6A, glm interaction: *β* = -6.11, *SE* =1.37, *z* = -4.46, *P* = 1.31 x 10^-4^, Table 3, third row) indicating significant differences in slope between the saline selection line (Line iii) and saline control line at 15 ppt (Line iv). Thus, only for *NHA-5* was the increase in expression over generations in the selection line significantly greater than the increase in the control line, reflecting the evolution of increased expression for *NHA-5* in the saline selection line and a divergence from the ancestral saline state at 15 ppt. The remaining 16 genes showed no significant expression changes and no consistent directional trends (increase or decrease) from Generation 1 to Generation 6 (S6 Fig; S5 Table). Expression levels of these genes remained relatively stable across generations in both the selection and control lines.

**Fig 6.**
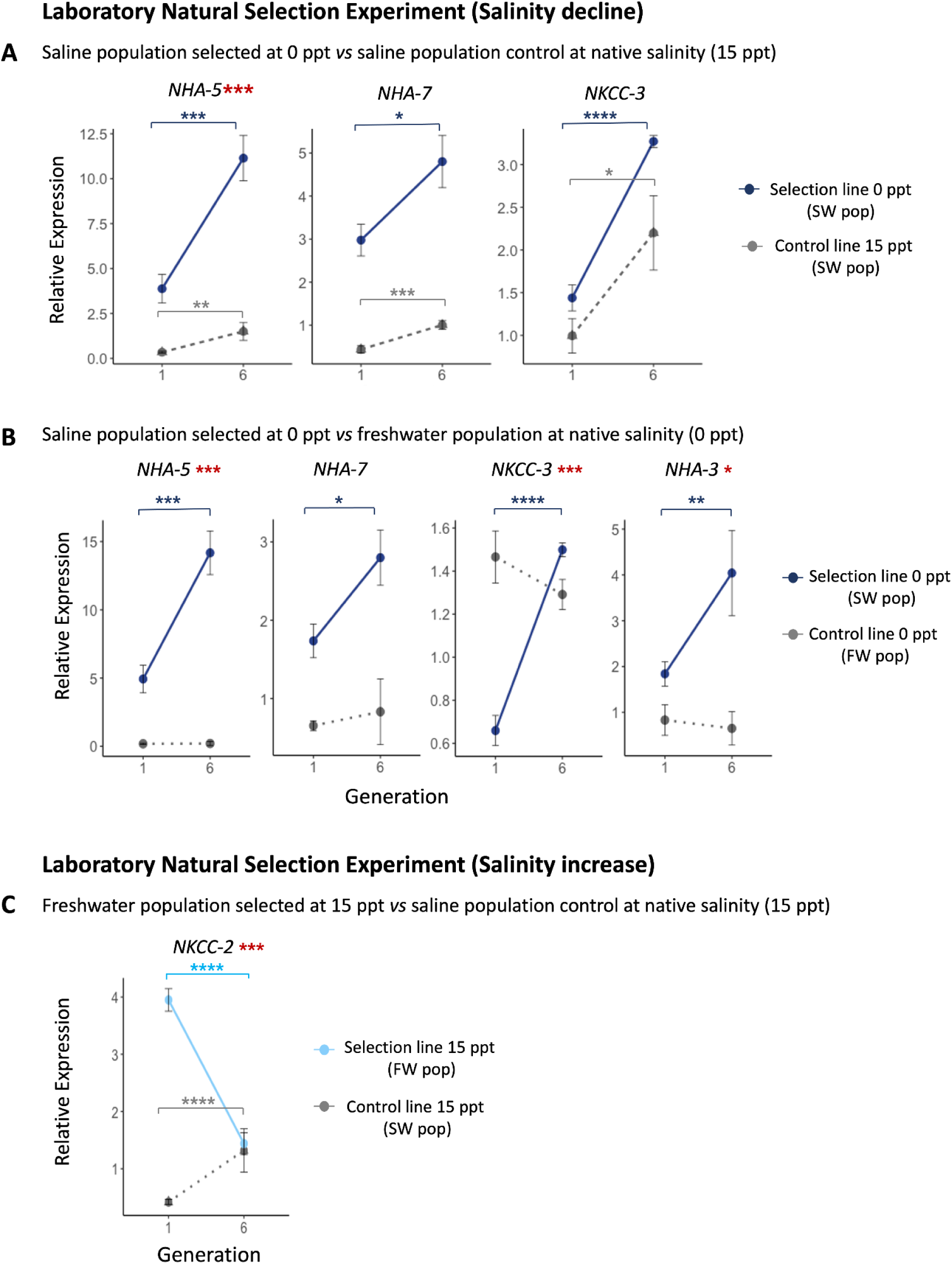
Ion transporter genes showing evolutionary shifts in expression following six generations of laboratory selection in response to salinity change. **(A)** Gene expression trajectories of selected ion transport-related genes in a saline population exposed to salinity decline (0 ppt) for six generations (Fig 2B, Line iii), relative to a saline control line maintained at native salinity (15 ppt) (Fig 2B, Line iv). Gene expression profiles for additional genes are shown in S6 Fig and full statistical results are shown in S5 Table. **(B)** Gene expression trajectories of selected ion transport-related genes in the same saline population undergoing salinity decline (0 ppt) for six generations (Fig 2B, Line iii), relative to a freshwater population control line maintained at native salinity (0 ppt) (Fig 2B, Line i). Results for additional genes are shown in S7 Fig and statistical results are presented in S6 Table. **(C)** Gene expression trajectory of *NKCC-2* in a freshwater population exposed to salinity increase (15 ppt) for six generations (Fig 2B, Line ii), relative to the saline ancestral population maintained at native salinity (15 ppt) (Fig 2B, Line iv). Full results are presented in S9 Fig and statistical results are shown in S8 Table. **“**Relative expression” here refers to the amount of ion transporter (and CA) mRNA expression relative to the mean expression of housekeeping genes (*EF-1α* and *GDPH*) (Equation 1 in Methods). Data points represent Mean ± SE. Dotted gray lines indicate the relevant native-salinity control for each comparison. Red **\***asterisks near gene names denote significant *Line x Generation* interaction effects (i.e., significantly different trajectory slopes etween the selection and control lines) based on generalized linear models (Table 3). Navy or blue asterisks **\***within trajectories indicate significant changes in expression from Generation 1 to Generation 6 based on linear models (Table 3). Level of significance indicated by asterisks (\**P* < 0.05, \*\**P* < 0.01, \*\*\**P* < 0.001, \*\*\*\**P* < 0.0001).

Additionally, for the salinity decline selection line at 0 ppt (Fig 2B, Line iii) in comparison with the freshwater population control maintained at 0 ppt (Fig 2B, Line i), four genes exhibited significant evolutionary shifts in gene expression. After six generations at 0 ppt, in the saline population selection line (Line iii) the genes *NHA-3*, *NHA-5*, *NHA-7* and *NKCC-3* increased in expression from Generation 1 to Generation 6 (Fig 6B, Table 3). In contrast, the ion transport-related genes in the freshwater population control line at 0 ppt (Line i) showed no significant change in expression from Generation 1 to Generation 6 (Figs 6B and S7; Tables 3 and S7). When the slopes of the saline selection line and the freshwater control line were compared to each other, the genes *NHA-5, NKCC-3,* and *NHA-3* exhibited significant *Line* x *Generation* interaction effects, indicating the divergence of slopes between the two lines. For NHA-3 and NHA-5, the gene expression pattern for the salinity decline selection line increased beyond that of the freshwater control (pattern of divergence) (glm interaction *NHA-3*: *β* = -2.39, *SE* = 0.90, *z* = -2.67, *P* = 0.0163 ; *NHA-5*: *β* = -9.23, *SE* = 2.27, *z* = -4.06, *P* = 6.70 x 10^-4^), while for *NKCC-3,* the saline selection line converged towards the freshwater control line (glm interaction: *β* = -1.01, *SE* = 0.209, *z* = -4.86, *P* = 1.49 x 10^-4^) (Fig 6B; Table 3). For *NHA-7*, the slopes of the selection and control lines did not significantly differ from each other (Fig 6B, Table 3). *NHA-1* also showed a significant increase in the saline selection line from Generation 1 to Generation 6, but did not diverge significantly from the freshwater control line (S7 Fig). The remaining 14 ion transport-related genes (S7 Fig) showed no significant expression change and no consistent directional trends, with expression remaining relatively stable in both the selection and control lines from Generation 1 to Generation 6 (S6 Table).

In the laboratory natural selection experiment for salinity increase, the freshwater population selected at 15 ppt for six generations (Fig 2B, Line ii) was compared to the freshwater population control line maintained at 0 ppt (Fig 2B, Line i) to assess divergence in gene expression. No genes showed significant expression changes between the selection and control lines (S8 Fig; S7 Table). However, when the freshwater selection line at 15 ppt (Line ii) was compared to the saline control at its native salinity (15 ppt) (Fig 2B, Line iv), *NKCC-2* was the only gene that exhibited a significant evolutionary shift, with expression in the selection line decreasing from Generation 1 to Generation 6 and converging toward the expression level of the saline control line (Fig 6C; Table 3). The significant convergence was reflected in the significant *Line* x *Generation* interaction effect (glm interaction: *β* = -3.62, *SE* = 0.418, *z* = -8.64, *P* = 1.60 x 10^-8^) (Table 3). Although several other ion transport-related genes also showed significant *Line* x *Generation* interaction effects, including NHA-5, NHA-6, NHA-7, and NKCC-3 (S9 Fig, S8 Table), these interactions were not driven by evolutionary change in the selection line, but rather by shifts in expression between Generation 1 and Generation 6 in the saline control line.

## DISCUSSION

Rapid responses to environmental change often involve physiological acclimation (plasticity), heritable genetic shifts (evolution), and/or their combined effects. In the *E. affinis* species complex, rapid adaptation to salinity decline has occurred repeatedly over the past ∼80 years, as estuarine populations have repeatedly colonized freshwater habitats across the Northern Hemisphere (5,7,29,34,36,40). During these rapid salinity transitions, ion transport-related gene families constitute the largest functional category of genes showing signatures of selection and/or evolutionary shifts in gene expression (21,29,36–38). Our study investigated the acclimatory and evolutionary shifts in ion transporters (and carbonic anhydrase, *CA*) gene expression during this salinity transition, focusing on the key genes that have previously shown the greatest and most consistent evolutionary shifts (21,29).

We quantified acclimatory and evolutionary shifts in ion transport-related gene expression by comparing ancestral saline and invasive freshwater populations across salinities in a common garden reaction norm experiment (Fig 2A). We also examined evolutionary shifts in gene expression in a laboratory natural selection experiment that imposed salinity change (Fig 2B). Within the expanded ion transport-related gene families of *E. affinis* complex, our results disentangled which paralogs showed evolutionary shifts in gene expression between populations and during laboratory selection in response to salinity change, and which showed plastic shifts in expression across salinities. Together, our results suggest that rapid responses to salinity change occur through coordinated acclimatory and/or evolutionary changes in ion transporter (and *CA*) paralog expression, allowing ion regulation to be reconfigured over remarkably short timescales.

### 1. Paralog-specific evolutionary and acclimatory responses in gene expression

The most striking pattern to emerge from the common garden experiment (Fig 2A) was that evolutionary and acclimatory shifts in gene expression in response to salinity change involved divergent patterns among different paralogs of the same gene families (*Hypothesis 4 in introduction*). These divergent patterns were evident across the massively expanded *NKA*, *NKCC*, *NHA*, and *CA* gene families in the *E. affinis* species complex. While other arthropods typically contain 1-2 ion transporter paralogs, *E. affini*s complex genomes are notable in containing 7-8 *NHA* paralogs, 8 *NKA*-*α* paralogs, 6 *NKCC* paralogs, and 13 *CA* paralogs (21,36,37,39,40,43). Our results suggest that these gene family expansions have potentially enabled subfunctionalization or neofunctionalization (56,57), where different paralogs may have become specialized for different aspects of iono/osmoregulatory function.

Within these gene families, we observed differences in the relative levels of evolutionary and acclimatory responses among paralogs. Some ion transport-related paralogs showed predominantly evolutionary divergence in gene expression between populations (e.g. *NKCC-3*, *NHA-5*), whereas others displayed primarily acclimatory responses across salinitines (e.g. *NKCC-2, NHA-4, CA-9*). Several ion transporters exhibited both evolutionary and acclimatory responses (e.g. *NKA-α-1*, *NHA-5*, *NHA-7*) (Fig 4). Notably, a subset of these paralogs (*NKA-α-1, NKCC-2, CA-9, NHA-5*, and *NHA-7)* also exhibited a significant *Population* x *Salinity* interaction, indicating that the plasticity of expression itself has evolved between the saline and freshwater populations (Fig 4). Together, these paralog-specific expression patterns are consistent with the hypothesis that gene family expansions have facilitated functional specialization across salinities, potentially supporting their adaptive responses at different salinities.

#### 1.1 Evolutionary responses in gene expression

We hypothesized that certain ion transport-related genes would show evolutionary shifts in expression between saline and freshwater populations (*Hypothesis 1*) and during laboratory selection in response to salinity change (*Hypothesis 6*). In support of *Hypothesis 1*, *NKA-α-1* and *NKCC-3* showed significantly higher expression in freshwater populations across the treatment salinities (Fig 4, Table 2). The elevated expression of *NKA-α-1* in freshwater populations across salinities (Fig 4 and 5) suggests that this paralog has properties that support ion uptake under ion-poor conditions, consistent with the higher hyperosmoregulatory capacity of the freshwater populations of *E. carolleeae* and *E. gulfia* relative to their ancestral saline populations (20).

This pattern of increased expression of *NKA-α-1* in the freshwater population was consistent with the previous genome-wide gene expression study (29), which similarly identified *NKA-α-1* as the only NKA paralog showing evolutionary increases in expression in freshwater relative to saline lines at both 0 and 15 ppt. However, overall enzyme activity of NKA was previously found to be higher in the saline populations than in the freshwater populations of *E. carolleeae* and *E. gulfia* under common garden conditions (23). In addition, *in situ* expression of NKA in the maxillary gland of *E. carolleeae* was found to be significantly elevated in the saline population, relative to the freshwater population, under common garden freshwater conditions (30). Thus, *NKA-α-1* appears to be an outlier among NKA paralogs in showing an evolution of increased expression in the freshwater population, relative to the saline population.

This stark difference in expression patterns of *NKA-α-1* relative to other paralogs (*NKA-a-2,* S4 Fig) (29) and relative to overall NKA expression and activity (23,30) suggests the specialized contribution of this NKA paralog to freshwater adaptation. The consistently elevated expression of *NKA-α-1* in the freshwater population in our study likely represents a key mechanism for performing ion uptake and maintaining hyperosmoregulatory capacity in freshwater environments. Na^+^/K^+^-ATPase establishes the electrochemical gradients necessary for secondary active ion transport and is highly expressed in osmoregulatory epithelia across taxa (58,59), including in copepod ionocytes in swimming legs, maxillary glands, and midgut (30,31,41,42).

Interestingly, across salinities, the freshwater population expressed higher levels of *NKA-α-1*, relative to *NKA-α-2*, whereas the saline population expressed greater levels of *NKA-α-2* than *NKA-α-1* (Fig 5). This pattern of *NKA* “paralog switching” that we observe between the saline and freshwater populations (Fig 5) is analogous to the NKA paralog switching that occurs within individual fish that migrate across salinities. In several teleost fish, such as salmonids (60,61), alewife (62), and zebrafish (63), alternative NKA *α*-subunit paralogs are expressed within individual fish as they migrate between habitat salinities. Notably, landlocked alewives show a dampened switching response relative to their diadromous ancestors (62), indicating that the switching response itself can evolve between populations.

The functional consequences of paralog switching in fish arise from a small number of amino acid substitutions in the transmembrane (TM) regions of the *α*-subunit that alter ion-binding and pump stoichiometry. In zebrafish, substitutions in TM5 and TM8 of the *α*_1a_.5 paralog block the third Na⁺ binding site, shifting stoichiometry from 3 Na⁺:2 K⁺:1 ATP to 1 Na⁺:1 K⁺:1 ATP and enabling osmoregulation in ion-poor freshwater (64). Additional TM-region substitutions modifying cation affinity and electrogenicity in other taxa have also been found (65). It is therefore possible that the population-level switching we report here shares similar structural differences between *NKA-α-1* and *NKA-α-2*, and future work should test whether the freshwater-favored paralog harbors substitutions in these same TM regions.

Divergence in expression patterns among paralogs was also evident in the *NKCC* gene family. The paralog *NKCC-3* showed clear evolution of increased expression in the freshwater population across all treatment salinities, relative to the saline population (Fig 4). On the other hand, *NKCC-2* showed an acclimatory shift toward elevated expression under freshwater conditions for both saline and freshwater populations and significantly greater expression in the freshwater population at higher salinity (Fig 4). This pattern suggests that these genes contribute to freshwater adaptation or acclimation. *NKCC-2* also showed a strong acclimatory increase at lower salinity (Fig 4; see next section *1.2* *Acclimarity Response*), suggesting that this paralog also contributes to freshwater acclimation. These results are consistent with the previous genome-wide RNA-seq study (29), which similarly found evolutionary upregulation of *NKCC-3* and *NKCC-2* in freshwater relative to saline inbred lines of the *E. affinis* complex, but only at 15 ppt, as well as a strong plastic upregulation of *NKCC-2* in both saline and freshwater lines at low salinity. Notably, this study also reported the opposite pattern for *NKCC-1* and *NKCC-4,* which were evolutionarily downregulated in the freshwater population relative to the saline population across salinities (29). These results strongly suggest that paralogs within the NKCC family have diverged in their functions at different salinities and contribute differentially to freshwater adaptation. Our results suggest that the paralogs *NKCC-2* and *NKCC-3* are likely the principal candidates for ion uptake in dilute environments and likely contribute to freshwater adaptation.

NKCC function has not been characterized in copepods. In other animals, the Na^+^/K^+^/2Cl^-^ cotransporter (NKCC) mediates the electroneutral, coupled transport of Na^+^, K^+^, and Cl^-^ across epithelial membranes, and members of this family are expressed in the osmoregulatory organs of fishes such as Atlantic Salmon (*Salmo salar*) and Tilapia, and of decapod crustaceans such as the sesarmid crab *Episesarma mederi*, including in enterocytes, renal tubular cells, and gill ionocytes (22,28,66,67). The direction of transport depends on the membrane domain in which a given paralog is expressed, for example: NKCCs localized in apical membranes of epithelial cells generally mediate ion absorption from the extracellular media (e.g. ambient water, intestinal lumen, excretory organ’s lumen), whereas basolaterally located NKCCs primarily function in ion secretion or intracellular ion recycling (68–70). This pattern is well documented in vertebrates, in which NKCC-1 (not homologous with *E. affinis* complex NKCC-1) is typically expressed in basolateral membranes (e.g. in gill ionocytes of seawater teleosts) while NKCC-2 (not homologous with *E. affinis* complex NKCC-2) is apical (e.g. in enterocytes and renal tubular cells) (71).

However, the role of NKCC in the *E. affinis* complex, and in copepods more generally, remains unresolved. No functional or localization study exists for any NKCC paralog in this system: neither the membrane domain (apical *versus* basolateral) nor the transport kinetics, activity, and regulation of these paralogs have been characterized. Consequently, the direction of ion movement and the physiological role of individual NKCC paralogs in *E. affinis* cannot yet be inferred from expression data alone.

Both *NHA-5* and *NHA-7* showed significant evolutionary shifts in expression between populations at 0 ppt, supporting *Hypothesis 1*. However, these shifts were in the opposite direction from what we had predicted, with higher expression in the saline population than in the freshwater population (Fig 4). This pattern is particularly notable for *NHA-7*, which had previously been identified as a key ionoregulatory gene showing evolutionary increases in expression in the freshwater population relative to the saline population under freshwater conditions (29). Several differences between the two studies could contribute to this discrepancy, including differences in primer specificity between the qPCR assays used here and the RNA-seq approach of Posavi et al. (29).

Tissue specific expression patterns appear to be consistent with the pattern found in Posavi et al. (29). Most notably, *in situ* immunolocalization had shown that NHA-7 was consistently expressed at much higher levels in the gut of freshwater population of *E. carolleeae* at native salinity (0 ppt), relative to the freshwater population at 15 ppt or the saline population at both 0 and 15 ppt (31). The midgut localization further raises the possibility that NHA-7 contributes to ion uptake from food under low-salinity conditions, since algal cells maintain an internal osmolarity equivalent to ∼2-5 ppt (72). However this interpretation remains speculative as the transport direction, membrane physiology, and precise role of NHA-7 in the *E. affinis* species complex have not been resolved. By contrast, NHA-5 has been localized in intracellular structures of the ionocytes in the swimming legs (Crusalis organs) (31). This localization suggests a different function related to intracellular pH regulation or vesicular ion transport within ionocytes, rather than direct transepithelial Na^+^ uptake.

Divergence in expression was also evident among the carbonic anhydrase (CA) paralogs (S4 Fig). Specifically, *CA-11* and *CA-12* showed higher expression in the saline population across all salinities (S4 Fig), consistent with the previous genome-wide RNA-Seq study (29), which reported CA paralogs 11, 12, 8, 7, and 14 as downregulated in freshwater lines relative to saline lines across salinities. However, in contrast to that study, we did not observe *CA-9* upregulation in the freshwater population at 0 ppt. Instead, *CA-9* showed increased expression at 15 ppt in both populations (see next section *1.2* *Acclimatory Response*). Carbonic anhydrases support acid-base balance by catalyzing the hydration of CO_2_ to generate H^+^ and HCO ^−^, key acid and base equivalents. These ions are subsequently exchanged for Na^+^ and Cl^-^ in osmoregulatory epithelia, facilitating the uptake of these ions (22,73,74). Thus, CAs serve as important enzymes in secondary active ion transport in ion-poor habitats (75,76).

Interestingly, several of the genes that showed evolutionary divergence in expression between the saline and freshwater populations in the common garden experiment (Fig 2A) also showed rapid evolutionary shifts in expression during laboratory natural selection (*Hypotheses 5* and *6*) (Fig 6). In particular, in the saline population undergoing salinity decline (Fig 2B, Line iii), *NHA-5*, *NHA-7*, and *NKCC-3* exhibited significant increases in expression after six generations at 0 ppt (Fig 6A, Table 3). However, only *NHA-5* exhibited significant change in slope relative to the control line at 15 ppt (Fig 2B, Lines iv;. The significant *Line* x *Generation* interaction (Table 3) indicates the significant deviation in trajectories between the selection and control lines. Interestingly, the evolution of increased expression of *NHA-5* at 0 ppt (Fig 6A) trended in the opposite direction of the native freshwater population, which showed reduced expression of these genes under freshwater (0 ppt) conditions (Fig 4). It is possible that the evolution of increased expression of *NHA-5* represents the initial evolutionary response, but that longer term freshwater adaptation might result in the reduction of expression due to other compensatory evolutionary changes.

We predicted that this saline selection line (Fig 2B, Line iii) would converge toward expression resembling the freshwater control line at 0 ppt (Fig 2B, Line i) after six generations of selection at 0 ppt. This convergence was evident only for *NKCC-3,* which showed increased expression in the saline selection line that converged towards the higher expression observed in the freshwater population control line (Fig 6B). The shift was consistent with the higher expression of *NKCC-3* in the freshwater population at 0 ppt, relative to the saline population (Fig 4). In contrast, expression of *NHA-5*, *NHA-7,* and *NHA-3* in the saline selection line (Fig 2B, Line iii) did not converge toward the freshwater control line at 0 ppt (Fig 2B, Line i) (Fig 6B). Instead, these genes exhibited increased expression under declining salinity (Fig 6B). This pattern was in stark contrast with the lower expression of *NHA-5* and *NHA-7* in the freshwater population in the common garden experiment (Fig 4). Again, for *NHA-5* and *NHA-7* this result suggests that the short term evolutionary response in the selection line differs from the long term evolutionary differences between the saline and freshwater populations.

Interestingly, in the freshwater selection line subjected to salinity increase (Fig 2B, Line ii), only *NKCC-2* converged toward the expression pattern of the saline control population (Fig 2B, Line iv) (Fig 6C). That is, *NKCC-2* showed a significant decline in expression after 6 generations at 15 ppt, such that its expression level converged with that of the saline control line at 15 ppt. This pattern was consistent with the reduced expression of *NKCC-2* in the saline population relative to the freshwater population at 15 ppt in the common garden comparison (Fig 4). Thus, in this case, short term selection did appear to recapitulate the evolutionary differences between the natural saline and freshwater populations.

Thus, for *NKCC-3* under salinity decline (Fig 6B) and *NKCC-2* under salinity increase, (Fig 6C) short-term evolution appeared to recapitulate the long-term expression profile of naturally saline-or freshwater-adapted populations (Fig 4), but for *NHA-5*, *NHA-7*, and *NHA-3*, it did not. Instead, for the *NHA* paralogs adaptation to salinity decline may involve divergent evolutionary trajectories on different time scales. The elevated expression of *NHA-5*, *NHA-7,* and *NHA-3* in the saline selection line subjected to salinity decrease (Fig 6A, 6B) may represent a response that provides short-term benefits, even if this expression pattern is later refined over longer evolutionary timescales (77,78).

Overall, other than *NKCC-2*, the freshwater selection line (Fig 2A, Line ii) subjected to salinity increase showed little evolutionary change in ion transport-related gene expression after six generations (Fig 6C; S8 Table). The limited response of the freshwater selection line may reflect differences in evolutionary history between the populations. The Lake Michigan freshwater population experiences consistently low salinity (300 *µ*S/cm), whereas the St. Lawrence saline population inhabits a seasonally fluctuating salinity regime (5 to 40 ppt) (35). Such environmental fluctuations in the ancestral saline habitat could maintain greater standing genetic variation through balancing selection, thereby increasing the capacity for rapid evolutionary response during salinity change (7,36,79–81). This difference in selection regime and standing genetic variation may help explain why the saline population showed a greater selection response to salinity change than the freshwater population (this pattern is explored more in section *2**. The Evolution of Plasticity* below).

#### 1.2 Acclimatory responses in gene expression

Our common garden experiment also revealed significant acclimatory responses (phenotypic plasticity) in gene expression across treatment salinities in both saline and freshwater populations (*Hypothesis 2*) (Figs 3, 4, and S4). Both saline and freshwater populations exhibited elevated expression of *NHA-5*, *NHA-7*, and *NKCC-2* at 0 ppt relative to 15 ppt (Fig 4, Table 1). *NHA-7* showed substantial upregulation at 0 ppt *vs* 15 ppt in the saline population and a more modest upregulation in the freshwater population (Fig 4, Table 1). For *NHA-5*, acclimatory upregulation at 0 ppt was observed only in the saline population (approximately 10-fold relative to 15 ppt), whereas expression in the freshwater population remained low at all salinities (Fig 4). This pattern for is consistent with the previous genome-wide study (29), in which *NHA-5* showed strong acclimatory upregulation at low salinity in the saline lines, weak and inconsistent plasticity in the freshwater lines, and a significant positive correlation between plastic responses in the saline lines and evolutionary shifts between populations. Thus, for *NHA-5*, the saline population maintained plasticity in gene expression across salinities, while the freshwater population evolved reduced plasticity (canalization) for this gene.

*NKCC-2* showed dramatic acclimatory responses with much greater fold change between 0 and 15 ppt in the saline population relative to the freshwater population (Table 1).Thus, although *NKCC-2* was clearly salinity responsive, its specific functional role cannot yet be inferred, given the absence of localization and functional data for the NKCC family in copepods (see section 1.1).

Both populations showed significantly elevated *CA-9* expression at 15 ppt compared to 0 ppt (Fig 4, Table 3). This result differed from the previous genome-wide RNA-Seq study (29), in which *CA-9* was upregulated in the freshwater population. This difference suggests that the environmental or physiological conditions regulating *CA-9* expression remain incompletely understood. There are 13 paralogs of CA in the *E. affinis* complex genome (43)), and expression patterns differ widely among them (S4 Fig) (29). Consequently, the elevated expression of *CA-9* at 15 ppt observed in our study is unlikely to reflect a role in ion secretion, because CA-generated H^+^ and HCO ^-^ are exchanged for Na^+^ and Cl^-^ mainly under low-salinity conditions (22,75). Instead, this pattern may indicate that *CA-9* has a role in acid-base balance or other physiological adjustments associated with more saline conditions. Salinity shifts can alter acid-base homeostasis in crustaceans, suggesting that CA paralogs expressed at higher salinities could contribute to pH regulation, CO_2_ handling, or other aspects of osmoconforming physiology (82). Future studies examining the functional roles and environmental regulation of individual CA paralogs would be worthwhile to pursue.

### 2. The evolution of plasticity in gene expression

Consistent with our *Hypothesis 3*, we found significant evolutionary shifts in plasticity in gene expression between saline and freshwater populations. We found evidence for the evolution of plasticity for expression of five ion transport-related genes (Fig 4), as indicated by the significant *Population* × *Salinity* interactions (S3 Table). This significant interaction effect indicated differences in the slope of the plastic responses between the populations, indicating the evolution of plastic responses to differences in salinity (83,84).

Both *NHA-5* and *NHA-7* exhibited significantly greater plasticity in expression (steeper reaction norm slopes) in the ancestral saline population compared to the invasive freshwater population (Fig 4). This result indicates that freshwater populations have evolved canalized (reduced) plasticity in expression for these genes relative to their saline ancestor (84,85). Canalization likely reflects loss of plasticity in stable environments, making plastic responses less necessary or even costly to maintain (54,83). The freshwater populations in Lake Michigan experience low salinity with minimal variation, whereas the saline populations in the St. Lawrence estuary experience seasonal fluctuations from 5 to 40 ppt (29). The negative genetic correlations and apparent tradeoffs between high and low salinity tolerance in *E. affinis* complex populations (9,86,87) would tend to lead to the loss of high salinity tolerance in a constant freshwater environment (88,89).

Interestingly, not all genes showed canalization in the freshwater population. *CA-9* showed marginally greater plasticity in freshwater populations than in saline populations, while *NKA-α-1* showed greater plasticity in the freshwater population, but with only a modest expression difference between 0 and 15 ppt (Fig 4) (Table 1).

Thus, our results indicate that plasticity itself is a trait that can evolve between saline and freshwater populations (Fig 4). Plastic responses are themselves adaptive traits, and in some cases plasticity alone may be sufficient for a population to persist in a novel environment, particularly when the plastic response is well matched to the new conditions and the environmental change is not extreme (88,89). In other cases, when plasticity is not sufficient, and long-term adaptation additionally requires heritable change in gene expression, including genetic variation in plasticity itself, for populations to track sustained environmental change (54,89). Determining the potential for future evolution of plasticity will require studies that examine the level of genetic variation in plasticity within populations, such as by examining clutch (genotype) specific reaction norms.

### 3. Applying our results to the predicted ion transport model

Our previous ion transport model for *E. affinis* complex had proposed that ion uptake in freshwater-adapted populations occurs through coordinated transporter activity in the osmoregulatory epithelia of the swimming legs, maxillary gland, and gut (21,22,30,31,41,42). In this model, CA supplies H^+^ ions that are pumped out of the cell by VHA, generating an electrochemical gradient across the apical membrane, while NKA pumps Na^+^ into the hemolymph and helps maintain the ionic conditions that favor Na^+^ uptake. NHA is then hypothesized to transport Na^+^ from the external medium into the cell. Our whole-organism gene expression results largely support this current model but indicate that the model should be refined to incorporate both paralog-specific functions of NKA, NHA, and CA. Our results here also point to the potential importance of NKCC, which was not included in the original crustacean model because its localization in *E. affinis* complex, or in any copepod, remains unknown. Because our measurements were made on whole animals rather than isolated tissues, these data identify candidate paralogs involved in osmoregulation but cannot assign their functions to particular organs or membrane domains (20,22,31).

The strongest support for the previous *E. affinis* complex model comes from the NKA results. The dramatic upregulation of *NKA-α-1* in the freshwater population across salinities (Fig 4) supports the prediction that enhanced NKA activity by *NKA-α-1* is essential for freshwater adaptation. In contrast, the lack of a comparable response in *NKA-α-2* suggests functional adaptive specialization among paralogs in different salinity environments rather than uniform regulation across the NKA gene family. Previous studies reported higher overall NKA activity and/or protein expression in saline populations under low-salinity conditions, but those studies did not distinguish among NKA paralogs (23,30). Thus, the apparent discrepancy between those results and our qPCR data likely reflects the fact that total NKA activity or protein abundance integrates multiple isoforms and tissues, whereas our analysis resolves individual paralogs but not their localization. Together, these results suggest that *NKA-α-1* may be the paralog more directly associated with adaptation in the freshwater population for ion uptake, even though total NKA activity or expression can remain higher in saline populations in some tissues.

Likewise, the *NHA* results (Fig 4; S4 Fig) indicate that the previous *E. affinis* ion-transport model should be revised to incorporate paralog-specific *NHA* paralogs rather than treating *NHA* as a single transport component. *NHA-7* fits the current model most closely, as it was upregulated at 0 ppt in both populations and has been localized to the gut (31), consistent with a role in Na^+^ uptake under low-salinity conditions. By contrast, both *NHA-7* and *NHA-5* were more highly expressed at 0 ppt in the saline population than in the freshwater population, showing that *NHA* expression patterns cannot be interpreted solely as signatures of freshwater adaptation. Previous localization data further indicate that these paralogs are localized in different cellular and tissue compartments, with NHA-7 detected in the gut and swimming legs and NHA-5 in intracellular structures of swimming-leg ionocytes (31). Thus, the model should differentiate among NHA paralogs explicitly (Fig 7), while future functional studies will be necessary to determine how these differences in expression and localization translate into distinct paralog-specific roles in ion transport.

**Fig 7.**
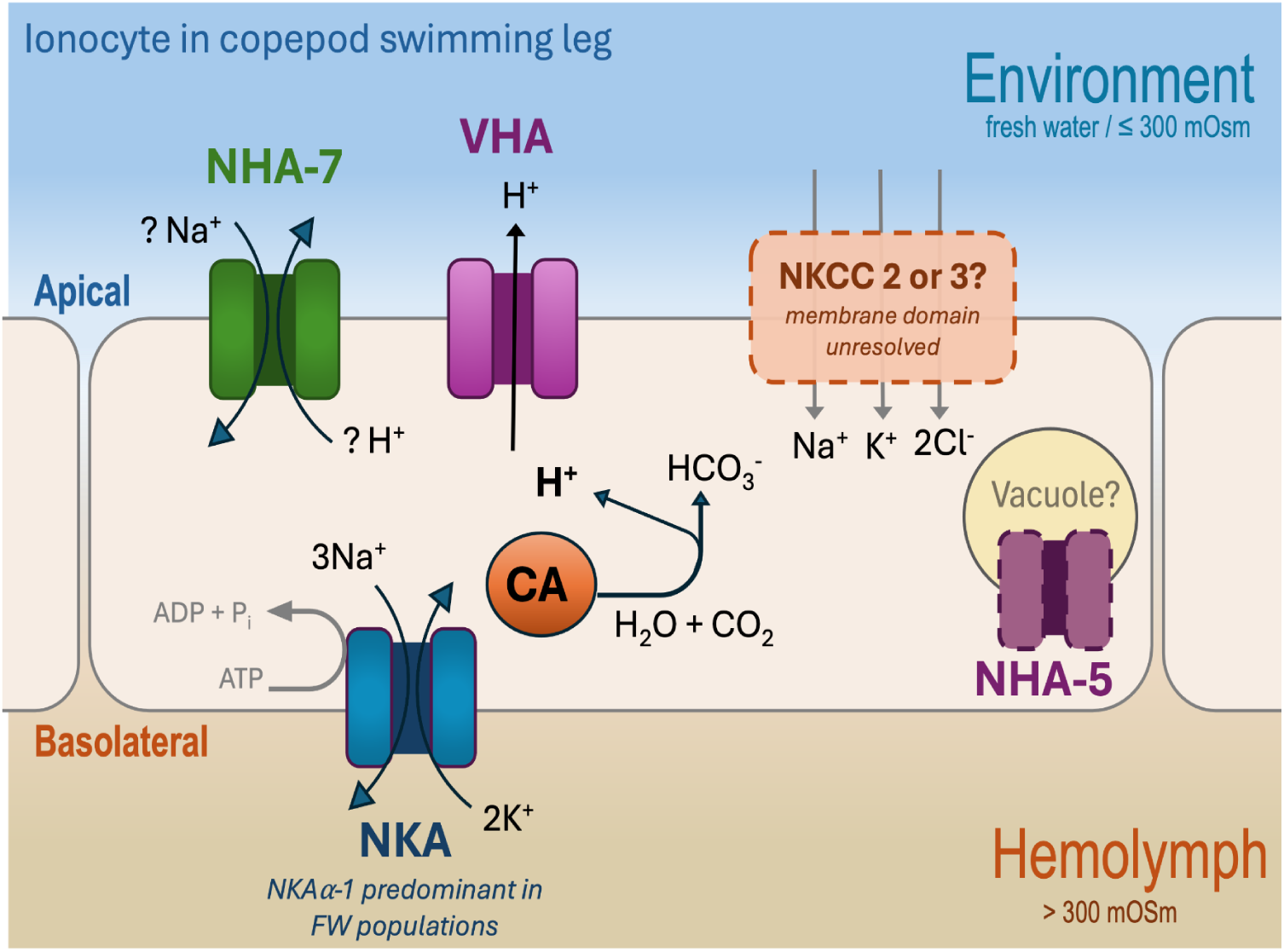
Theoretical model of ion transport in the osmoregulatory ionocytes of a swimming leg in *E. carolleeae* under freshwater conditions, revised to incorporate paralog-specific findings. The model shows VHA pumping H^+^ out across the apical membrane and generating the electrochemical gradient that energizes ion uptake, while CA supplies the H ^+^ and HCO_3_^-^ for this exchange. Among the NHA paralogs, NHA-7 best fits the original uptake model, as it is upregulated at 0 ppt in both populations and localized to the apical membrane in the gut and swimming legs (Popp 2024). NHA-5 localizes to intracellular structures (e.g. vacuoles) in the cytoplasm, with its function unresolved. NKCC (paralog 2 or 3) is included as a candidate route for coupled Na^+^, K^+^, and Cl^-^ transport. NKCC was absent from the original crustacean model because the membrane domain has not been determined for any NKCC paralog in copepods. At the basolateral membrane, NKA maintains the gradient that drives apical Na^+^ entry, with NKAα-1 predominating in freshwater populations. The model depicts freshwater conditions only, and salinity-specific differences in expression are reported in Figs 4 to 6.

Although *NKCC* was not included in the original crustacean ion-uptake model because its localization within *E. affinis* complex tissues remains unknown, our results suggest that NKCC should now be considered in future updates of the model (Fig 7). *NKCC-3* showed evolutionary upregulation in the freshwater population across salinities, whereas *NKCC-2* showed stronger acclimatory upregulation under freshwater conditions, indicating distinct paralog responses to salinity (Fig 4). Together with previous evidence that *NKCC-1* and *NKCC-4* are upregulated in saline relative to freshwater lines (29), these results support functional divergence within the NKCC family. However, direct tissue-specific and subcellular localization studies will be required before specific NKCC paralogs can be assigned roles in apical ion uptake, basolateral ion recycling, or ion secretion.

The patterns of expression of *CA* and *VHA* (Fig 4, S4 Fig) suggest that the functional roles of these gene families also require more intricate refinement in the model. The acclimatory upregulation of *CA-9* at 15 ppt (Fig 4) and the evolutionary upregulation of *CA-11* and *CA-12* in saline populations (S4 Fig) are consistent with a role for carbonic anhydrases in acid-base regulation and in supplying H^+^ and HCO ^−^ for ion transport. By contrast, the two VHA subunits (*VHA-H* and *VHA-a*) examined showed relatively limited evolutionary or acclimatory shifts in expression, despite high overall expression (Fig 5 and S4, Table 2). This pattern of *VHA* expression may reflect its multiple cellular functions beyond osmoregulation, including roles in vesicular trafficking, protein degradation, and intracellular pH regulation (26,90).

Overall, our results emphasize the importance of recognizing the different functions of ion transporter and CA paralogs. Thus, refinement of the *E. affinis* complex ion-transport model will depend on gaining more information on the localization and functional specialization among the paralogs (56,57). Rather than treating "NKA," "NKCC," or "NHA" as single functional units, our enhanced model should incorporate paralog-specific, tissue-specific, and ultimately membrane-specific roles that will be revealed by and tested directly in future localization and functional studies.

## METHODS

### 1. Population sampling

Population sampling was performed in 2020 for both saline and freshwater populations of *Eurytemora carolleeae* (Atlantic clade of the *Eurytemora affinis* complex) (34,35,51,52). The saline population was sampled in October of 2020 from a tidal marsh in the St. Lawrence estuary, in Baie de L’isle Verte, Quebec, Canada (48°00’14” N, 69°25’31” W), where salinity fluctuates seasonally between 5 – 40 ppt. Individuals from a freshwater population were collected in summer of 2020 from Lake Michigan at Racine harbor, Wisconsin, USA (42°43′46″N, 87°46′44″W), where salinity is relatively constant at 0 ppt (conductivity of 300 *µ*S/cm) throughout the year.

### 2. Experiment 1: Common garden reaction norm experiment

A common garden reaction norm experiment was performed to determine the evolutionary shifts in ion transporter gene expression between ancestral saline and freshwater invading populations of *E. carolleeae* from the St. Lawrence drainage system, as well as the evolution of plasticity between the populations and plasticity in expression across salinities within each population (Fig 2A). Both populations were first reared under a common intermediate salinity (5 ppt) to remove the effects of prior environmental acclimation to their native salinities. Then, the populations were reared at a range of salinities (0, 5, and 15 ppt) to disentangle genetic (evolutionary) and environmental (plastic) responses to gene expression (see details below).

Prior to the experiment, both saline and freshwater populations were maintained in the laboratory for two years (approximately 24 generations) at their native salinities (15 and 0 ppt respectively; photoperiod 15L:9D). Saline populations were fed the saltwater alga *Rhodomonas salina*, and freshwater populations were fed the freshwater alga *Rhodomonas minuta*. Freshwater populations were cultured in fresh water obtained from the Hérault river, France, and saline populations were maintained in a mix of fresh water and natural seawater obtained from Sète, France (SMEL, Station Méditerranéenne de l’Environnement Littoral, Université Montpellier).

The common garden experiment was conducted in 2 blocks at the Marine Biodiversity, Exploitation and Conservation (MARBEC) laboratory in Montpellier, France. The first block was performed in July 2022, whereas the second block was conducted in September 2022. At the start of each experimental block, 60 ovigerous females from the saline population (St. Lawrence estuary, Canada) and 60 from the freshwater population (Lake Michigan, Wisconsin, USA) were transferred to a common salinity of 5 ppt. Once the nauplii (larvae) hatched, adult females were removed from the beakers and the nauplii were reared under this common salinity (5 ppt) until metamorphosing to the copepodid (juvenile) stage (Fig 2A). Groups of approximately 200 juveniles were then gradually transferred to treatment salinities of 0, 5, and 15 ppt, using either a salinity decline (5 → 2.5 → 1 → 0 ppt) or a salinity increase (5 → 8 → 10 → 15 ppt) regime with groups transferred to a new salinity every two days. Each population–salinity treatment was replicated in six biological replicates per block (12 biological replicates [beakers] across both blocks) (N = 72 total treatments across populations and salinities). These treatment salinities (0, 5, 15 ppt) represent conditions proximate to those in their native habitat for *E. carolleeae* populations, where 0 ppt (lake water) corresponds to the native freshwater habitat, 15 ppt to the native saline habitat, and 5 ppt serves as an intermediate control well tolerated by both populations (86,87). These juveniles then matured into adulthood at the treatment salinities.

To avoid imposing selection on early life stages, the populations were not maintained at the treatment salinities for more than one generation. Previous studies have shown that 0 and 15 ppt induce high mortality and impose strong selection on the larval stages that are not native to those salinities(9,86). Additionally, maternal effects associated with rearing salinity have been shown to exert minimal influence on offspring ion transporter (VHA and NKA) enzyme activity (23). After one week of exposure to the treatment salinities as adults, groups of 50 copepods were preserved for gene expression analysis in RNAlater for 24h at 4°C and were then stored at -20°C until mRNA extraction.

### 3. Experiment 2: Laboratory natural selection experiment

Continuing from the common garden experiment (Fig 2A), a laboratory natural selection experiment was conducted to uncover evolutionary shifts in ion transporter gene expression in response to rapid salinity change under controlled laboratory conditions (Fig 2B). At the end of the common garden experiment, after adults had been sampled for gene expression analysis, the remaining animals were consolidated from the replicate beakers into a single beaker for each population x salinity treatment (0 and 15 ppt; i.e., a single beaker per treatment; Fig 2B, lines i–iv). Approximately 100 individuals remained in each replicate beaker after sampling, such that each consolidated beaker initially contained approximately 600 animals at Generation 1. The salinity change protocol was that of the common garden experiment: animals were brought to the treatment salinity step-wise within a single generation (decline: 5 → 2.5 → 1 → 0 ppt; increase: 5 → 8 → 10 → 15 ppt, with transfer to a new salinity every two days; see *Experiment 1*), and each beaker was then maintained at its treatment salinity for an additional six generations. Comparisons were made between the following lines to determine differences in gene expression:

1. *<u>Salinity Decline</u>* <u>experiment</u>: the saline population selection line, reared at 0 ppt for six generations (Fig 2B, line *iii*) was compared to two types of controls:

- (Control 1) <u>Saline population maintained at its native salinity</u> (15 ppt; Fig 2B, line *iv*), enabling the comparison between the saline population response to declining salinity *vs.* the control saline population with no selection.
- (Control 2) <u>Freshwater population maintained at its native salinity</u> (0 ppt; Fig 2B, line *i*), enabling the comparison between the saline population selected for freshwater tolerance (0 ppt) *vs.* a natural freshwater population at 0 ppt.
2. *<u>Salinity Increase</u>* <u>experiment</u>: the freshwater population selection line, reared at 15 ppt for six generations (Fig 2B, line *ii*), was compared to two types of controls:

- (Control 1) <u>Freshwater population maintained at its native salinity</u> (0 ppt; Fig 2B, line *i*), enabling the comparison between the freshwater population response to increasing salinity *vs.* the control freshwater population with no selection.
- (Control 2) <u>Saline population maintained at its native salinity</u> (15 ppt; Fig 2B, line *iv*), enabling the comparison between the freshwater population selected for high salinity tolerance (15 ppt) *vs.* a natural saline population at 15 ppt.

After six generations at the treatment salinity, adults were collected for gene expression analysis. In this experiment, a *replicate* was defined as a pool of 50 animals taken from a single beaker. There were a total of 4 beakers, one per population-salinity treatment (Fig 2B-lines *i - iv*), to increase power, multiple independent pools were collected from each beaker. In total, 20 independent pools were generated across selection and control lines. The final number of replicates varied depending on survival: saline selection line (Fig 2B - line *iii*: N = 4), saline control line (Fig 2B - line *iv:* N = 6), freshwater selection line (Fig 2B - line *ii:* N = 6), and freshwater control line (Fig 2B - line *i:* N = 4).

The animals were preserved in RNAlater for 24 hours at 4°C and then moved to -20°C until RNA extraction. Our goals were to determine: (1) the extent to which selection lines evolve in response to salinity change and (2) whether six generations of selection at 0 or 15 ppt results in shifts in gene expression towards the pattern of native populations. To address these goals, mRNA expression levels of key iono/osmoregulatory genes were compared across selection lines and control populations described above.

### 4. RNA extraction and reverse transcription

RNA extraction was performed for 50 adult copepods (pools of 25 males + 25 females) from the common garden and selection experiments (Fig 2). An RNeasy Micro Kit (Qiagen, Germany; Ref: 74104) was used for total RNA extraction and DNAse treatment. The A260/A280 and A260/A230 ratios were measured using a NanoDrop® One Spectrophotometer (ThermoFisher, USA). RNA was only used when the A260/A280 ratio was above 1.9 and A260/A230 ratio was above 2.0. First-strand cDNA synthesis was conducted with 800 ng of RNA using qScript™ cDNA SuperMix (Quanta Biosciences™) and the necessary reagents including random primers, oligo primers, and qScript reverse transcriptase. The resulting cDNA samples were stored at -20°C until qPCR was performed.

### 5. Primer design and Polymorphism in the primer binding regions

Specific primer pairs were designed for selected iono/osmoregulatory gene paralogs of interest and for the housekeeping genes using Geneious Prime® 2023.0.1.The primers used in this study for both the saline and freshwater populations are indicated in Table 4. The presence of polymorphic sites between the saline and freshwater populations within each primer’s binding region was investigated to ensure that differences in expression levels between populations were not biased by primer binding differences. Using a two step approach, the *E. carolleeae* reference genome (39) was scanned for single nucleotide polymorphisms (SNPs) between saline and freshwater populations. First, the coding sequence of each ion transporter was aligned to the repeat-masked reference genome using BLAST. The command line version of BLAST (v.2.12.0) was used, and the reference genome was formatted and indexed as a BLAST-searchable reference using the *-makeblastdb* utility. The *-blastn* option was employed to run BLAST for each gene’s coding sequence. The coordinates of each gene in the reference genome from the BLAST outputs with identities > 95% were gathered.

**Table 4:**
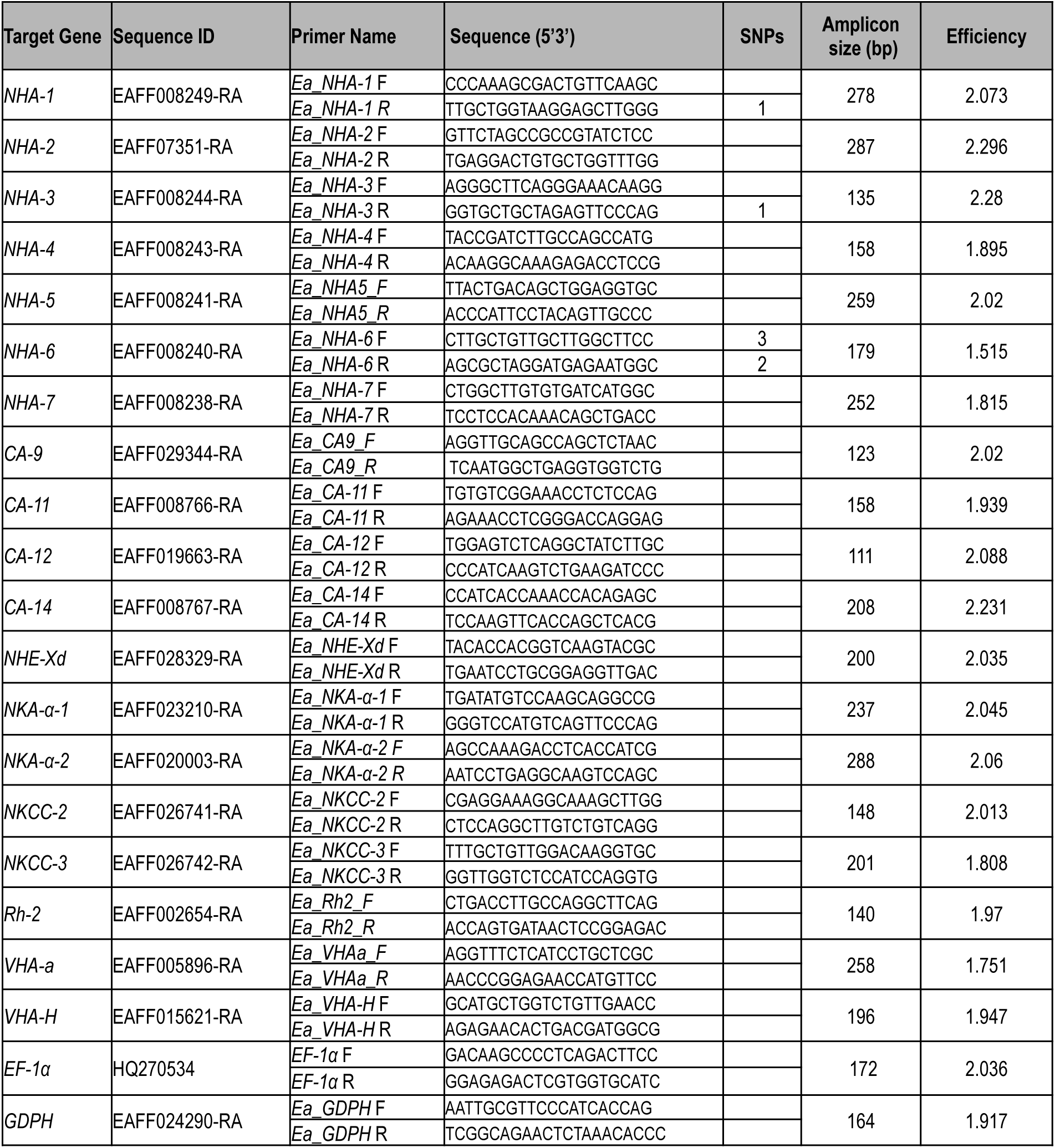
List of forward (F) and reverse (R) primers designed for 19 ion transport-related genes and 2 housekeeping genes from the *E. carolleeae* genome for qRT-PCR analysis. The table also indicates the amplicon size (bp), efficiency of the primer pairs, and the presence of SNPs with the primer sequence between saline and freshwater populations.

Second, primer pairs were aligned to the coding sequence of their corresponding gene using AliView (v. 1.28;(91)) in order to obtain the position of each primer on the coding sequence of each gene. The primer’s positions on the reference genome were deduced by calculating the distance between a primer’s position on the coding sequence and the start of that coding sequence on the reference genome. The primers’ regions were scanned for SNPs using previously published whole-genome Pool-seq data from wild saline and freshwater populations (36). Briefly, processed reads generated by Illumina HiSeq sequencing were mapped to the repeat-masked *E. carolleeae* genome using BWA-MEM (v.0.7.17;(92)) (see (36) for details on sequencing strategy and raw read processing). Duplicated reads were removed using Sambamba *-markdup* option (v.0.8.0; (93)) and variant calling was performed using FreeBayes (v.1.3.5;(94)). Finally, the VCF file containing all SNPs in R (v.4.3.1; (95)) was imported using the *poolfstats* package (v.2.2.0; (96)) and data were filtered for SNPs with a minimum allele frequency < 0.10.

The primers used in this study for both the saline and freshwater populations are indicated in Table 4, along with the presence and number of SNPs within the primer sequences. For genes with primers that did not include SNPs, both evolutionary and acclimatory comparisons between saline and freshwater populations across salinities were conducted. For genes with SNPs in the priming sequences between saline and freshwater populations, the populations were not compared to each other; for these genes, comparisons were performed across salinities within each population to determine plastic responses to salinity for that population.

### 6. Quantitative real time RT-PCR

To determine evolutionary and acclimatory shifts in ion transporter gene expression, quantitative RT-PCR was performed for 19 ion transporter genes in saline and freshwater populations across three salinities (0, 5, 15 ppt). Standard curves were generated with a mix of all samples to determine the optimal cDNA dilution for the analyzed genes, as well as the primer pair efficiency (Table 4). The qRT-PCR cocktail mix included 0.75 *μ*L of SensiFAST™ SYBR® No-ROX Kit (Bioline, UK), each primer at 0.4 *μ*M concentration, 0.19 *μ*L of ultra-pure water, and 0.5 *μ*L of diluted cDNA (diluted at 1/16). The samples with qRT-PCR cocktail were dispensed into 384-well reaction plates in an Echo®525 liquid handling system (Labcyte Inc., San Jose, CA, USA). qRT-PCR was performed in duplicates for each sample using a real-time PCR thermal cycler (LightCycler 480, Roche). The qRT-PCR profiles consisted of an initial denaturation step (1 cycle, 95 °C, 150 s), followed by denaturation (45 cycles, 95 °C, 15 s), annealing (1 cycle, 60 °C, 5 s), elongation (1 cycle, 72 °C, 10 s) and a final step (1 cycle, 40 °C, 30 s). The amplification specificity was controlled by a melting curve program. The no-template control used ultra-pure water in this program.

Relative amount of mRNA expression between populations, salinities, or selection lines was determined for each ion transport-related gene, accounting for primer efficiency (E) and the expression levels of internal controls. The relative gene expression was calculated as the ratio between the threshold cycle (*Ct*) of the target genes and the geometric mean of two housekeeping genes (97) elongation factor-1 alpha (*EF-1α*) (98) and glyceraldehyde 3-phosphate dehydrogenase (*GDPH*). The calculation of relative expression also accounted for PCR primer binding efficiencies (E) for each gene and the difference in threshold cycle (*ΔCt)* of an internal control (see next paragraph) and the sample (99). The threshold cycle (*Ct*) is the cycle number at which the fluorescence from the amplifying DNA exceeds a predefined threshold, indicating a significant increase in signal intensity. The mean *Ct* of both housekeeping genes was evaluated to verify that it did not vary across experimental conditions, and it was confirmed to remain consistent between populations, treatment salinities, and between Generation 1 and Generation 6 in the laboratory natural selection experiment (*P* < 0.05). The relative amount of ion transporter mRNA expression (*Relative Expression*) between populations or salinities was calculated as:

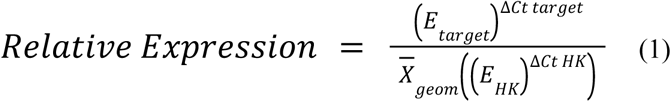

where *E* = efficiency of PCR primer, *target* = gene of interest, *HK* = housekeeping gene, *ΔCt target* = difference between *Ct* of the internal control (see next paragraph) and the individual sample for the target gene, and *ΔCt HK* = difference between *Ct* of the housekeeping gene’s internal control condition and the individual sample for the housekeeping gene.

The internal control remained consistent within each comparison between conditions but varied among experiments depending on the type of analysis. In the common garden reaction norm comparisons (examining the effects of *Population*, *Salinity*, and *Population* × *Salinity*), the ancestral saline population at 5 ppt was used as the internal control for genes without SNP differences in the primer pair between populations. If SNPs were present in the primer regions, gene expression levels were not compared between populations; instead, expression levels within each population were normalized to the 5 ppt treatment of each population. When comparing expression levels of different ion transporter genes to one another, the mean expression of *NHA-7* was used as the internal control, separately for the saline and freshwater populations. *NHA-7* was chosen as this reference because it was a top target of selection and showed robust expression responses to salinity change in previous studies of the *E. affinis* complex (29,36), making it a biologically meaningful anchor for comparison. Additionally, using a single, common internal control placed the expression of all ion transport-related genes on the same scale (Fig 5), enabling pairwise comparisons among NHA paralogs and among all genes. For the Principal Component Analysis (PCA), the average expression of all genes at 5 ppt was used as the internal control to evaluate relative gene expression trends.

In the laboratory natural selection experiment, changes in gene expression after six generations of natural selection (salinity increase or salinity decline) was observed in the selection lines *vs.* the control lines. For all the comparisons (see section *Experiment 2: Laboratory Natural Selection Experiment*), relative gene expression was calculated using the mean *Ct* level between the comparisons (described above) at Generation 1 as the internal control. This method enabled us to assess how the trajectory of gene expression patterns evolved following laboratory selection in response to salinity increase or salinity decline.

### 7. Statistical analyses

All statistical analyses and figure preparation were performed using R version 4.4.0 (95). First, a Principal Component Analysis (PCA) was employed to distinguish gene expression patterns of 15 ion transport-related genes between populations (saline *vs.* freshwater populations) and between salinity treatments (0 *vs.* 15 ppt) from the common garden experiment (Fig 2A). Key gene contributors to each principal component were identified, and the distribution of data points along the PC1 *vs.* PC2 and PC1 *vs.* PC3 were used to elucidate population-specific (ie. evolutionary shifts) and salinity-induced (ie. acclimatory shifts) differences in ion transporter gene expression patterns. A multivariate analysis of variance (Two-way MANOVA) was applied using the *PCA scores* as the dependent variable, and using the factors *Population* (freshwater *vs.* saline), *Salinity* (0 *vs.* 15 ppt), and their interaction *Population* x *Salinity*. This approach allowed for comparisons of gene expression across multiple axes of variation, capturing correlated expression changes not evident in univariate analyses.

Subsequently, for individual ion transport-related genes, the effects of fixed factors *Population* (saline *vs.* freshwater populations), *Salinity* (0, 5, 15 ppt), and *Population x Salinity* on gene expression levels (the response variable *Relative Expression,* Equation 1 above) from the common garden reaction norm experiment were analyzed using a two-way ANOVA. Initial analyses included *Block* as a fixed effect to account for potential variation in culturing conditions between blocked experiments. As *Block* effects were not significant for any gene, we proceeded with two-way ANOVA models with a pooled dataset. This two-way ANOVA was only conducted for the 16 genes for which the PCR primer sequences contained no SNPs between the saline and freshwater populations (Table 4). For the 3 ion transporter genes for which SNPs were detected in at least one of the primers (Table 4, *NHA-1, NHA-3, and NHA-6*), a one-way ANOVA was used to examine the effect of the factor *Salinity* on *Relative Expression* across salinities for each population. That is, as SNP differentiation in the PCR primers between the saline and freshwater populations could result in differential PCR priming and amplification between the populations, expression levels between the populations were not compared. However, examining the acclimatory responses across salinities within each population was possible. When the effects of factors were found to be significant, pairwise comparisons between treatments were performed using a *post hoc* Tukey’s HSD test.

Following the ANOVA analysis, additional tests were conducted to measure the magnitude of change in expression for the 19 iono/osmoregulatory genes (1) between the saline and freshwater populations and (2) across salinity treatments (0, 5, 15 ppt). To make these comparisons two metrics were calculated, the Log_2_(FC) in gene expression was calculated between each population at each treatment salinity and also within each population between treatment salinities.

In addition, for each population at each treatment salinity (i.e. freshwater population at 0 ppt, saline population at 0 ppt, freshwater population at 15 ppt, and saline population at 15 ppt), expression levels of different ion transport-related genes, as well as groups of genes, were compared to one another. Comparisons among groups of genes were conducted as follows: (1) between ion pumps (ATPases *NKA* and *VHA*) *vs.* all other ion transport-related genes, using a pooled Mann Whitney test, (2) between the 2 *NKA* paralogs (*NKA-α1* vs. *NKA-α2*), using a Student’s t-test, and (3) pairwise comparisons between members of the *NHA* gene family, using Student’s t-tests and accounting for multiple comparisons using a Bonferroni correction.

Finally, for the laboratory natural selection experiment, we examined the effects of selection on gene expression in two ways: first, by comparing changes between Generation 1 and Generation 6 within an *Experimental Line*, and second, by assessing differences in the trajectory of gene expression between the *Experimental Lines* (Fig 2, below the red line), by determining the significance of the *Line (Experimental Line) x Generation* interaction effect. Here, *Experimental Line* refers to the selection lines and the control lines explained in the above section: *Experiment 2: Laboratory natural selection experiment*.

First, to analyze the change in gene expression following six generations of natural selection, the Log_2_(FC) was calculated for each selection line or control line between Generations 1 and 6. In addition, Cohen’s *d* was computed as a standardized effect size, defined as the difference between group means divided by the pooled standard deviation. Second, a linear regression model (lm) was applied to examine the trajectory of gene expression between Generations 1 and 6, with *Generation* as the fixed factor and *Relative Expression* as the response variable. Finally, a generalized linear model (glm) was used to assess whether the trajectory of gene expression (slope from Generation 1 to Generation 6) differed between the selection and control lines (*Experimental Line)*. This model evaluated the *Line* x *Generation* interaction effect on the *Relative Expression* response variable, where a significant interaction effect indicates that the trajectory (slope) of gene expression differs between the two lines.

For both the common garden and natural selection experiments, the response variable of *Relative Expression* (Equation 1, above) was log transformed for each gene due to evidence of skewness and heteroscedasticity (S1 and S2 Figs).

## Data availability

All data supporting the results in this study are provided in the Supplementary Tables and Figures. The Illumina RNA-seq data compared in this study were previously published (29).

## Acknowledgements

This project was funded by National Science Foundation grants IOS-2412790, DEB-2055356, and OCE-1658517 and French National Research Agency ANR-19-MPGA-0004 (Macron’s “Make Our Planet Great Again” award) to Carol E. Lee. This project was also funded by NSF PRFB 2409904 to Teresa E. Popp.

## References

1. Crowl TA, Crist TO, Parmenter RR, Belovsky G, Lugo AE. The spread of invasive species and infectious disease as drivers of ecosystem change. Front Ecol Environ. 2008;6(5):238–46. doi:10.1890/070151

2. Simberloff D, Rejmánek M, editors. Encyclopedia of biological invasions. Berkeley (CA): University of California Press; 2011.

3. Lee CE, Downey K, Colby RS, Freire CA, Nichols S, Burgess MN, et al. Recognizing salinity threats in the climate crisis. Integr Comp Biol. 2022;62(2):441–60. doi:10.1093/icb/icac069

4. Williamson M, Fitter A. The varying success of invaders. Ecology. 1996;77:1661–6. doi:10.2307/2265769

5. Lee CE. Evolution of invasive populations. In: Simberloff D, Rejmánek M, editors. Encyclopedia of biological invasions. Berkeley (CA): University of California Press; 2011. p. 215–22.

6. Lee CE. Evolutionary genetics of invasive species. Trends Ecol Evol. 2002;17(8):389–95. doi:10.1016/S0169-5347(02)02554-5

7. Lee CE, Gelembiuk GW. Evolutionary origins of invasive populations. Evol Appl. 2008;1(3):427–48. doi:10.1111/j.1752-4571.2008.00039.x

8. Lee CE, Bell MA. Causes and consequences of recent freshwater invasions by saltwater animals. Trends Ecol Evol. 1999;14(7):284–8. doi:10.1016/S0169-5347(99)01596-7

9. Lee CE, Remfert JL, Gelembiuk GW. Evolution of physiological tolerance and performance during freshwater invasions. Integr Comp Biol. 2003;43:439–49.

10. Casties I, Seebens H, Briski E. Importance of geographic origin for invasion success: a case study of the North and Baltic Seas versus the Great Lakes–St. Lawrence River region. Ecol Evol. 2016;6(22):8318–29. doi:10.1002/ece3.2528

11. Holeck KT, Mills EL, MacIsaac HJ, Dochoda MR, Colautti RI, Ricciardi A. Bridging troubled waters: biological invasions, transoceanic shipping, and the Laurentian Great Lakes. BioScience. 2004;54(10):919–29. doi:10.1641/0006-3568(2004)054[0919:BTWBIT]2.0.CO;2

12. Ricciardi A. Patterns of invasion in the Laurentian Great Lakes in relation to changes in vector activity. Divers Distrib. 2006;12(4):425–33. doi:10.1111/j.1366-9516.2006.00262.x

13. Spidle AP, Marsden JE, May B. Identification of the Great Lakes quagga mussel as *Dreissena bugensis* from the Dnieper River, Ukraine, on the basis of allozyme variation. Can J Fish Aquat Sci. 1994;51(7):1485–9. doi:10.1139/f94-148

14. Cristescu MEA, Hebert PDN, Witt JDS, MacIsaac HJ, Grigorovich IA. An invasion history for *Cercopagis pengoli* based on mitochondrial gene sequences. Limnol Oceanogr. 2001;46(2):224–9. doi:10.4319/lo.2001.46.2.0224

15. Gelembiuk GW, May GE, Lee CE. Phylogeography and systematics of zebra mussels and related species. Mol Ecol. 2006;15(4):1033–50. doi:10.1111/j.1365-294X.2006.02816.x

16. May GE, Gelembiuk GW, Panov VE, Orlova MI, Lee CE. Molecular ecology of zebra mussel invasions. Mol Ecol. 2006;15(4):1021–31. doi:10.1111/j.1365-294X.2006.02814.x

17. Remane A, Schlieper C. Biology of brackish water. 2nd ed. New York (NY): John Wiley and Sons; 1971. p. 1–210.

18. Khlebovich VV, Abramova EN. Some problems of crustacean taxonomy related to the phenomenon of Horohalinicum. Hydrobiologia. 2000;417(1):109–13. doi:10.1023/A:1003863623267

19. Charmantier G, Charmantier-Daures M, Towle D. Osmotic and ionic regulation in aquatic arthropods. In: Evans DH, editor. Osmotic and ionic regulation: cells and animals. Boca Raton (FL): CRC Press; 2009. p. 165–320. doi:10.1201/9780849380525

20. Lee CE, Posavi M, Charmantier G. Rapid evolution of body fluid regulation following independent invasions into freshwater habitats. J Evol Biol. 2012;25(4):625–33. doi:10.1111/j.1420-9101.2012.02459.x

21. Lee CE. Ion transporter gene families as physiological targets of natural selection during salinity transitions in a copepod. Physiology. 2021;36(6):335–49. doi:10.1152/physiol.00009.2021

22. Lee CE, Charmantier G, Lorin-Nebel C. Mechanisms of Na⁺ uptake from freshwater habitats in animals. Front Physiol. 2022;13:1006113. doi:10.3389/fphys.2022.1006113

23. Lee CE, Kiergaard M, Gelembiuk GW, Eads BD, Posavi M. Pumping ions: rapid parallel evolution of ionic regulation following habitat invasions. Evolution. 2011;65(8):2229–44. doi:10.1111/j.1558-5646.2011.01308.x

24. Bourne SD, Hudson J, Holman LE, Rius M. Marine invasion genomics: revealing ecological and evolutionary consequences of biological invasions. In: Oleksiak MF, Rajora OP, editors. Population genomics: marine organisms. Cham (Switzerland): Springer; 2020. p. 363–98. doi:10.1007/13836_2018_21

25. Chintapalli VR, Kato A, Henderson L, Hirata T, Woods DJ, Overend G, et al. Transport proteins NHA1 and NHA2 are essential for survival, but have distinct transport modalities. Proc Natl Acad Sci U S A. 2015;112(37):11720–5. doi:10.1073/pnas.1508031112

26. Boudour-Boucheker N, Boulo V, Charmantier-Daures M, Anger K, Charmantier G, Lorin-Nebel C. Osmoregulation in larvae and juveniles of two recently separated *Macrobrachium* species: expression patterns of ion transporter genes. Comp Biochem Physiol A Mol Integr Physiol. 2016;195:39–45. doi:10.1016/j.cbpa.2016.02.005

27. Brennan RS, Healy TM, Bryant HJ, La V, Schulte PM, Whitehead A. Integrative population and physiological genomics reveals mechanisms of adaptation in killifish. Mol Biol Evol. 2018;35(11):2639–53. doi:10.1093/molbev/msy154

28. Boonsanit P, Chanchao C, Pairohakul S. Effects of hypo-osmotic shock on osmoregulatory responses and expression levels of selected ion transport-related genes in the sesarmid crab *Episesarma mederi* (H. Milne Edwards, 1853). Comp Biochem Physiol A Mol Integr Physiol. 2024;288:111541. doi:10.1016/j.cbpa.2023.111541

29. Posavi M, Gulisija D, Munro JB, Silva JC, Lee CE. Rapid evolution of genome-wide gene expression and plasticity during saline to freshwater invasions by the copepod *Eurytemora affinis* species complex. Mol Ecol. 2020;29(20):4835–56. doi:10.1111/mec.15681

30. Popp TE, Hermet S, Fredette-Roman J, McKeel E, Zozaya W, Baumlin C, et al. Evolution of ion transporter Na+/K+-ATPase expression in the osmoregulatory maxillary glands of an invasive copepod. iScience. 2024;27(7):110278. doi:10.1016/j.isci.2024.110278

31. Popp TE. Evolution of ion transporter expression during salinity decline [dissertation]. Madison (WI): University of Wisconsin–Madison; 2024. ProQuest publication no. 3093531525.

32. Lee CE. Evolutionary mechanisms of habitat invasions, using the copepod *Eurytemora affinis* as a model system. Evol Appl. 2016;9(1):248–70. doi:10.1111/eva.12334

33. Lee CE. Genome architecture underlying salinity adaptation in the invasive copepod *Eurytemora affinis* species complex: a review. iScience. 2023;26(10):107851. doi:10.1016/j.isci.2023.107851

34. Lee CE. Rapid and repeated invasions of fresh water by the copepod *Eurytemora affinis*. Evolution. 1999;53(5):1423–34. doi:10.2307/2640889

35. Winkler G, Dodson JJ, Lee CE. Heterogeneity within the native range: population genetic analyses of sympatric invasive and noninvasive clades of the freshwater invading copepod *Eurytemora affinis*. Mol Ecol. 2008;17(1):415–30. doi:10.1111/j.1365-294X.2007.03480.x

36. Stern DB, Lee CE. Evolutionary origins of genomic adaptations in an invasive copepod. Nat Ecol Evol. 2020;4(8):1084–94. doi:10.1038/s41559-020-1201-y

37. Stern DB, Anderson NW, Diaz JA, Lee CE. Genome-wide signatures of synergistic epistasis during parallel adaptation in a Baltic Sea copepod. Nat Commun. 2022;13:3610. doi:10.1038/s41467-022-31622-8

38. Diaz JA, Stern DB, Lee CE. Local adaptation despite gene flow in copepod populations across salinity and temperature gradients in the Baltic Sea and North Sea region. Divers Distrib. 2026;32(5):e70176. doi:10.1111/ddi.70176

39. Du Z, Gelembiuk G, Moss W, Tritt A, Lee CE. The genome architecture of the copepod *Eurytemora carolleeae* — the highly invasive Atlantic clade of the *Eurytemora affinis* species complex. Genomics Proteomics Bioinformatics. 2024;22(5). doi:10.1093/gpbjnl/qzae066

40. Du Z, Wirtz J, Zhou YJ, Jenstead A, Opgenorth T, Puls A, et al. Genome architecture evolution in an invasive copepod species complex. Nat Commun. 2025;16(1):10312. doi:10.1038/s41467-025-65292-z

41. Johnson KE, Perreau L, Charmantier G, Charmantier-Daures M, Lee CE. Without gills: localization of osmoregulatory function in the copepod *Eurytemora affinis*. Physiol Biochem Zool. 2014;87(2):310–24. doi:10.1086/674319

42. Gerber L, Lee CE, Grousset E, Blondeau-Bidet E, Boucheker NB, Lorin-Nebel C, et al. The legs have it: in situ expression of ion transporters V-type H⁺-ATPase and Na⁺/K⁺-ATPase in the osmoregulatory leg organs of the invading copepod *Eurytemora affinis*. Physiol Biochem Zool. 2016;89(3):233–50. doi:10.1086/686323

43. Zhou YJ, Lee CE. Evolutionary history of the α-carbonic anhydrase gene family in the phylum Arthropoda, with comparisons to αCA in chordates. Genome Biol Evol. 2026;18(7):evag156. doi:10.1093/gbe/evag156

44. Xiang MA, Linser PJ, Price DA, Harvey WR. Localization of two Na⁺- or K⁺-H⁺ antiporters, AgNHA1 and AgNHA2, in *Anopheles gambiae* larval Malpighian tubules and the functional expression of AgNHA2 in yeast. J Insect Physiol. 2012;58(4):570–9. doi:10.1016/j.jinsphys.2011.12.009

45. Kolosov D, Tauqir M, Rajaruban S, Piermarini PM, Donini A, O’Donnell MJ. Molecular mechanisms of bi-directional ion transport in the Malpighian tubules of a lepidopteran crop pest, *Trichoplusia ni*. J Insect Physiol. 2018;109:55–68. doi:10.1016/j.jinsphys.2018.06.005

46. Sterling KM, Harvey WR. Inhibition of sodium–hydrogen antiport by antibodies to NHA1 in brush border membrane vesicles from whole *Aedes aegypti* larvae. J Membr Biol. 2019;252(1):1–16. doi:10.1007/s00232-018-0053-8

47. Naseem MT, Beaven R, Koyama T, Naz S, Su SY, Leader DP, et al. NHA1 is a cation/proton antiporter essential for the water-conserving functions of the rectal complex in *Tribolium castaneum*. Proc Natl Acad Sci U S A. 2023;120(13):e2217084120. doi:10.1073/pnas.2217084120

48. Marioni JC, Mason CE, Mane SM, Stephens M, Gilad Y. RNA-seq: an assessment of technical reproducibility and comparison with gene expression arrays. Genome Res. 2008;18(9):1509–17. doi:10.1101/gr.079558.108

49. Everaert C, Luypaert M, Maag JLV, Cheng QX, Dinger ME, Hellemans J, et al. Benchmarking of RNA-sequencing analysis workflows using whole-transcriptome RT-qPCR expression data. Sci Rep. 2017;7:1559. doi:10.1038/s41598-017-01617-3

50. SEQC/MAQC-III Consortium. A comprehensive assessment of RNA-seq accuracy, reproducibility and information content by the Sequencing Quality Control Consortium. Nat Biotechnol. 2014;32(9):903–14. doi:10.1038/nbt.2957

51. Lee CE. Global phylogeography of a cryptic copepod species complex and reproductive isolation between genetically proximate “populations.” Evolution. 2000;54(6):2014–27. doi:10.1111/j.0014-3820.2000.tb01245.x

52. Alekseev VR, Souissi A. A new species within the *Eurytemora affinis* complex (Copepoda: Calanoida) from the Atlantic coast of USA, with observations on eight morphologically different European populations. Zootaxa. 2011;2767(1):41–56. doi:10.11646/zootaxa.2767.1.4

53. Waddington CH. Genetic assimilation of an acquired character. Evolution. 1953;7(2):118–26.

54. Lande R. Adaptation to an extraordinary environment by evolution of phenotypic plasticity and genetic assimilation. J Evol Biol. 2009;22(7):1435–46. doi:10.1111/j.1420-9101.2009.01754.x

55. Chevin LM, Lande R. When do adaptive plasticity and genetic evolution prevent extinction of a density-regulated population? Evolution. 2010;64(4):1143–50. doi:10.1111/j.1558-5646.2009.00875.x

56. Kuzmin E, Taylor JS, Boone C. Retention of duplicated genes in evolution. Trends Genet. 2022;38(1):59–72. doi:10.1016/j.tig.2021.06.016

57. Kondrashov FA, Rogozin IB, Wolf YI, Koonin EV. Selection in the evolution of gene duplications. Genome Biol. 2002;3(2):research0008.1. doi:10.1186/gb-2002-3-2-research0008

58. Thabet R, Ayadi H, Koken M, Leignel V. Homeostatic responses of crustaceans to salinity changes. Hydrobiologia. 2017;799(1):1–20. doi:10.1007/s10750-017-3232-1

59. Griffith MB. Toxicological perspective on the osmoregulation and ionoregulation physiology of major ions by freshwater animals: teleost fish, crustacea, aquatic insects, and Mollusca. Environ Toxicol Chem. 2017;36(3):576–600. doi:10.1002/etc.3676

60. Bystriansky JS, Richards JG, Schulte PM, Ballantyne JS. Reciprocal expression of gill Na⁺/K⁺-ATPase α-subunit isoforms α1a and α1b during seawater acclimation of three salmonid fishes that vary in their salinity tolerance. J Exp Biol. 2006;209(10):1848–58. doi:10.1242/jeb.02188

61. McCormick SD, Regish AM, Christensen AK, Björnsson BT. Differential regulation of sodium–potassium pump isoforms during smolt development and seawater exposure of Atlantic salmon. J Exp Biol. 2013;216(7):1142–51. doi:10.1242/jeb.080440

62. Colby RS, McCormick SD, Velotta JP, Jockusch E, Schultz ET. Paralog switching facilitates diadromy: ontogenetic, microevolutionary and macroevolutionary evidence. Oecologia. 2024;205(3):571–86. doi:10.1007/s00442-024-05588-x

63. Dymowska AK, Boyle D, Schultz AG, Goss GG. The role of acid-sensing ion channels in epithelial Na⁺ uptake in adult zebrafish (*Danio rerio*). J Exp Biol. 2015;218(8):1244–51. doi:10.1242/jeb.113118

64. Dymowska AK, Hwang PP, Goss GG. Structure and function of ionocytes in the freshwater fish gill. Respir Physiol Neurobiol. 2012;184(3):282–92. doi:10.1016/j.resp.2012.08.025

65. Jorgensen PL. Importance for absorption of Na⁺ from freshwater of lysine, valine and serine substitutions in the α1a-isoform of Na,K-ATPase in the gills of rainbow trout ( *Oncorhynchus mykiss*) and Atlantic salmon (*Salmo salar*). J Membr Biol. 2008;223(1):37–47. doi:10.1007/s00232-008-9111-y

66. Freire CA, Onken H, McNamara JC. A structure-function analysis of ion transport in crustacean gills and excretory organs. Comp Biochem Physiol A Mol Integr Physiol. 2008;151(3):272–304. doi:10.1016/j.cbpa.2007.05.008

67. Hiroi J, McCormick SD. New insights into gill ionocyte and ion transporter function in euryhaline and diadromous fish. Respir Physiol Neurobiol. 2012;184(3):257–68. doi:10.1016/j.resp.2012.07.019

68. Piermarini PM, Akuma DC, Crow JC, Jamil TL, Kerkhoff WG, Viel KCMF, et al. Differential expression of putative sodium-dependent cation-chloride cotransporters in *Aedes aegypti*. Comp Biochem Physiol A Mol Integr Physiol. 2017;214:40–9. doi:10.1016/j.cbpa.2017.09.007

69. Hoffmann EK, Schettino T, Marshall WS. The role of volume-sensitive ion transport systems in regulation of epithelial transport. Comp Biochem Physiol A Mol Integr Physiol. 2007;148(1):29–43. doi:10.1016/j.cbpa.2006.11.023

70. Castrop H, Schieß1 M. Physiology and pathophysiology of the renal Na-K-2Cl cotransporter (NKCC2). Am J Physiol Renal Physiol. 2014;307(9):F991–1104. doi:10.1152/ajprenal.00432.2014

71. Takvam M, Wood CM, Kryvi H, Nilsen T. Ion transporters and osmoregulation in the kidney of teleost fishes as a function of salinity. Front Physiol. 2021;12:664588. doi:10.3389/fphys.2021.664588

72. Raven JA. Costs and benefits of low intracellular osmolarity in cells of freshwater algae. Funct Ecol. 1995;9(5):701–7. doi:10.2307/2390241

73. Tseng YC, Yan JJ, Furukawa F, Chen RD, Lee JR, Tsou YL, et al. Teleostean fishes may have developed an efficient Na⁺ uptake for adaptation to the freshwater system. Front Physiol. 2022;13:947958. doi:10.3389/fphys.2022.947958

74. Krogh A. The active uptake of ions into cells and organisms. Proc Natl Acad Sci U S A. 1939;25(6):275–7. doi:10.1073/pnas.25.6.275

75. Gilmour KM, Perry SF. Carbonic anhydrase and acid–base regulation in fish. J Exp Biol. 2009;212(11):1647–61. doi:10.1242/jeb.029181

76. Henry RP, Lucu Č, Onken H, Weihrauch D. Multiple functions of the crustacean gill: osmotic/ionic regulation, acid-base balance, ammonia excretion, and bioaccumulation of toxic metals. Front Physiol. 2012;3:431. doi:10.3389/fphys.2012.00431

77. Stoebel DM, Hokamp K, Last MS, Dorman CJ. Compensatory evolution of gene regulation in response to stress by *Escherichia coli* lacking RpoS. PLoS Genet. 2009;5(10):e1000671. doi:10.1371/journal.pgen.1000671

78. Velotta JP, Cheviron ZA. Remodeling ancestral phenotypic plasticity in local adaptation: a new framework to explore the role of genetic compensation in the evolution of homeostasis. Integr Comp Biol. 2018;58(6):1098–110. doi:10.1093/icb/icy117

79. Posavi M, Gelembiuk GW, Larget B, Lee CE. Testing for beneficial reversal of dominance during salinity shifts in the invasive copepod *Eurytemora affinis*, and implications for the maintenance of genetic variation. Evolution. 2014;68(11):3166–83. doi:10.1111/evo.12502

80. Barrett RDH, Schluter D. Adaptation from standing genetic variation. Trends Ecol Evol. 2008;23(1):38–44. doi:10.1016/j.tree.2007.09.008

81. Lande R, Shannon S. The role of genetic variation in adaptation and population persistence in a changing environment. Evolution. 1996;50(1):434–7. doi:10.1111/j.1558-5646.1996.tb04504.x

82. Whiteley NM, Scott JL, Breeze SJ, McCann L. Effects of water salinity on acid–base balance in decapod crustaceans. J Exp Biol. 2001;204(5):1003–11. doi:10.1242/jeb.204.5.1003

83. Van Buskirk J, Steiner UK. The fitness costs of developmental canalization and plasticity. J Evol Biol. 2009;22(4):852–60. doi:10.1111/j.1420-9101.2009.01685.x

84. Flatt T. The evolutionary genetics of canalization. Q Rev Biol. 2005;80(3):287–316. doi:10.1086/432265

85. Shaw JR, Hampton TH, King BL, Whitehead A, Galvez F, Gross RH, et al. Natural selection canalizes expression variation of environmentally induced plasticity-enabling genes. Mol Biol Evol. 2014;31(11):3002–15. doi:10.1093/molbev/msu241

86. Lee CE, Petersen CH. Effects of developmental acclimation on adult salinity tolerance in the freshwater-invading copepod *Eurytemora affinis*. Physiol Biochem Zool. 2003;76(3):296–301. doi:10.1086/375433

87. Lee CE, Remfert JL, Chang YM. Response to selection and evolvability of invasive populations. Genetica. 2007;129(2):179–92. doi:10.1007/s10709-006-9013-9

88. Chevin LM, Lande R, Mace GM. Adaptation, plasticity, and extinction in a changing environment: towards a predictive theory. PLoS Biol. 2010;8(4):e1000357. doi:10.1371/journal.pbio.1000357

89. Chevin LM, Lande R. Adaptation to marginal habitats by evolution of increased phenotypic plasticity. J Evol Biol. 2011;24(7):1462–76. doi:10.1111/j.1420-9101.2011.02279.x

90. Tresguerres M. Novel and potential physiological roles of vacuolar-type H⁺-ATPase in marine organisms. J Exp Biol. 2016;219(14):2088–97. doi:10.1242/jeb.128389

91. Larsson A. AliView: a fast and lightweight alignment viewer and editor for large datasets. Bioinformatics. 2014;30(22):3276–8. doi:10.1093/bioinformatics/btu531

92. Li H, Durbin R. Fast and accurate short read alignment with Burrows-Wheeler transform. Bioinformatics. 2009;25(14):1754–60. doi:10.1093/bioinformatics/btp324

93. Tarasov A, Vilella AJ, Cuppen E, Nijman IJ, Prins P. Sambamba: fast processing of NGS alignment formats. Bioinformatics. 2015;31(12):2032–4. doi:10.1093/bioinformatics/btv098

94. Garrison E, Marth G. Haplotype-based variant detection from short-read sequencing [Internet]. arXiv; 2012 [cited 2024 Apr 27]. Available from: http://arxiv.org/abs/1207.3907

95. R Core Team. R: a language and environment for statistical computing. Vienna (Austria): R Foundation for Statistical Computing; 2020.

96. Gautier M, Vitalis R, Flori L, Estoup A. f-statistics estimation and admixture graph construction with Pool-Seq or allele count data using the R package poolfstat. Mol Ecol Resour. 2022;22(4):1394–416. doi:10.1111/1755-0998.13557

97. Vandesompele J, De Preter K, Pattyn F, Poppe B, Roy N, De Paepe A, et al. Accurate normalization of real-time quantitative RT-PCR data by geometric averaging of multiple internal control genes. Genome Biol. 2002;3(7):RESEARCH0034. doi:10.1186/gb-2002-3-7-research0034

98. Lauritano C, Borra M, Carotenuto Y, Biffali E, Miralto A, Procaccini G, et al. First molecular evidence of diatom effects in the copepod *Calanus helgolandicus*. J Exp Mar Biol Ecol. 2011;404(1–2):79–86. doi:10.1016/j.jembe.2011.05.009

99. Pfaffl MW. A new mathematical model for relative quantification in real-time RT-PCR. Nucleic Acids Res. 2001;29(9):e45. doi:10.1093/nar/29.9.e45

