## Supplementary Figures S1-S9 for "Evolutionary and acclimatory shifts in ion transporter gene expression across salinity transitions in the copepod *Eurytemora carolleeae*"

Romain GROS

Carol Eunmi LEE*
ORCID: 0000-0001-6355-0542

Catherine LORIN-NEBEL*
ORCID: 0000-0001-6966-0080

[1]Department of Biology, University of Wisconsin, 430 Lincoln Drive, Madison, WI 53706, USA

[2]MARBEC, Univ. Montpellier, CNRS, IRD, Ifremer, Montpellier, France

[†]Current address: Department of Entomology, The Ohio State University

Keywords: ion transporters, rapid evolution, acclimation, gene expression, natural selection experiment, common garden experiment

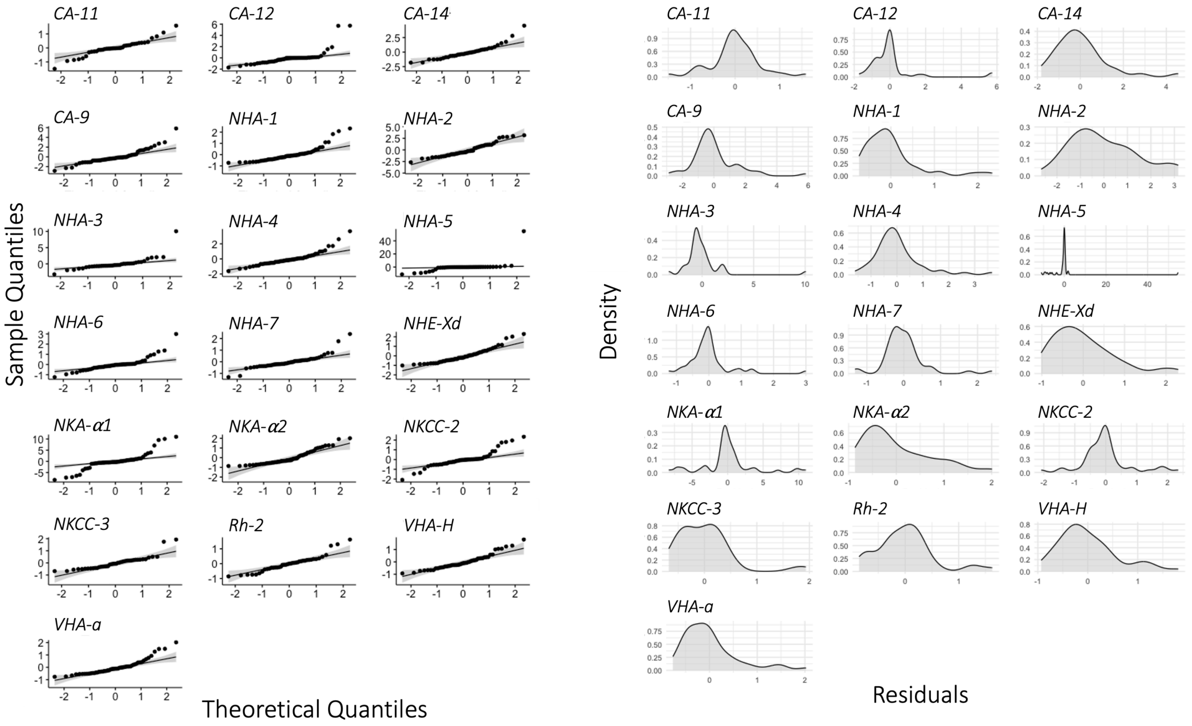


**Supplementary Figure S1. Normality assessment of the residuals for non-transformed *Relative Expression* of each gene across saline (SW) and freshwater (FW) populations at treatment salinities of 0, 5, and 15 ppt under common garden conditions. (A)** Quantile-quantile (Q-Q) plot displaying the distribution of empirical residuals against the theoretical quantiles. Points outside the confidence interval (gray shaded area) suggest that the residuals do not follow a normal distribution. **(B)** Density plot showing the distribution of residuals values (x-axis). The plot indicates an a-symmetrical distribution of residuals indicating evidence of skewness. *Relative Expression* values were subsequently log transformed (S2 Fig).

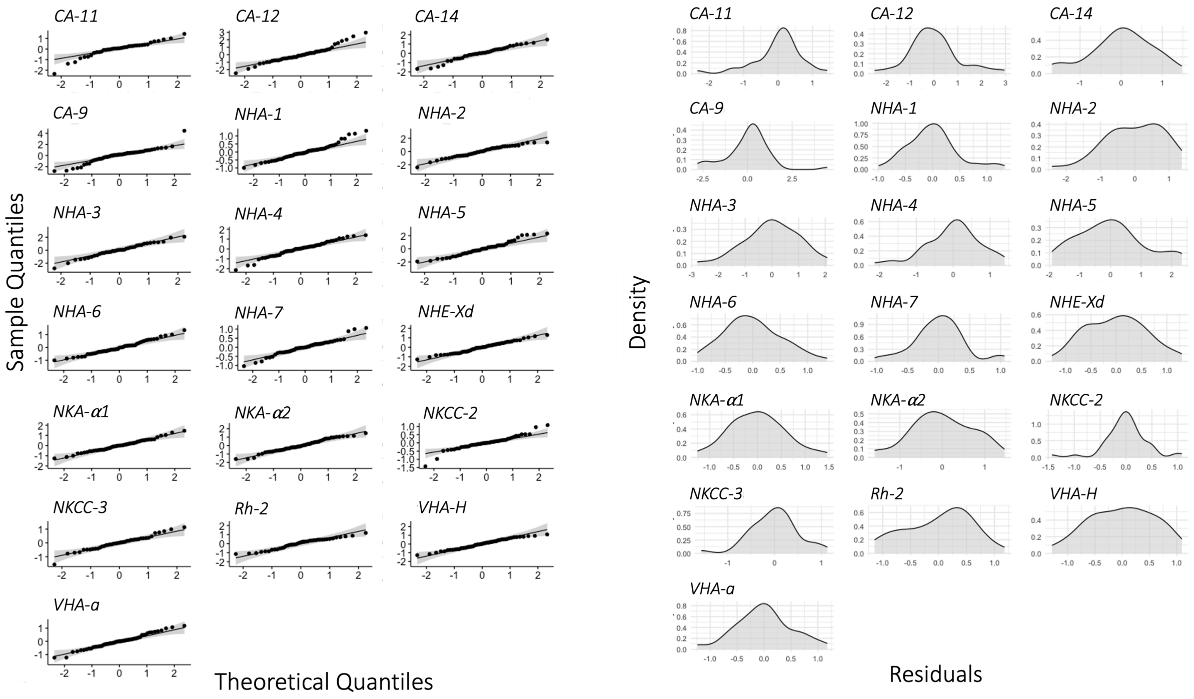


**Supplementary Figure S2. Normality assessment of the residuals for log-transformed *Relative Expression* of each gene across saline (SW) and freshwater (FW) populations at treatment salinities of 0, 5, and 15 ppt under common garden conditions. (A)** Quantile-quantile (Q-Q) plot displaying the distribution of empirical residuals against the theoretical quantiles. Points within the confidence interval (gray shaded area) suggest that the residuals follow a normal distribution. **(B)** Density plot showing the distribution of residuals values (x-axis). The plot indicates a symmetrical distribution of residuals with no strong evidence of skewness.

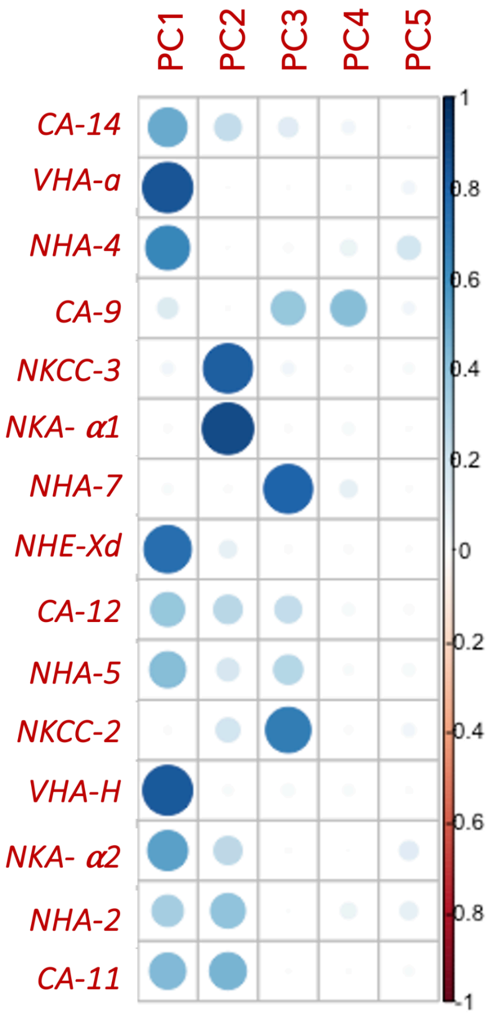


**Supplementary Figure S3. Contributions of the ion transport-related genes to multiple principal components (PCs) in the PCA analysis, based on gene expression levels from the common garden experiment.** Only genes without SNPs in the primers are shown. The analysis highlights the varying influence of different genes on each PC, reflecting the saline and freshwater population responses to salinity shifts (0 and 15 ppt) in the common garden experiment. The degree of each gene's contribution is depicted by the color and size of the circles, as indicated by the scale on the right side of the graph.

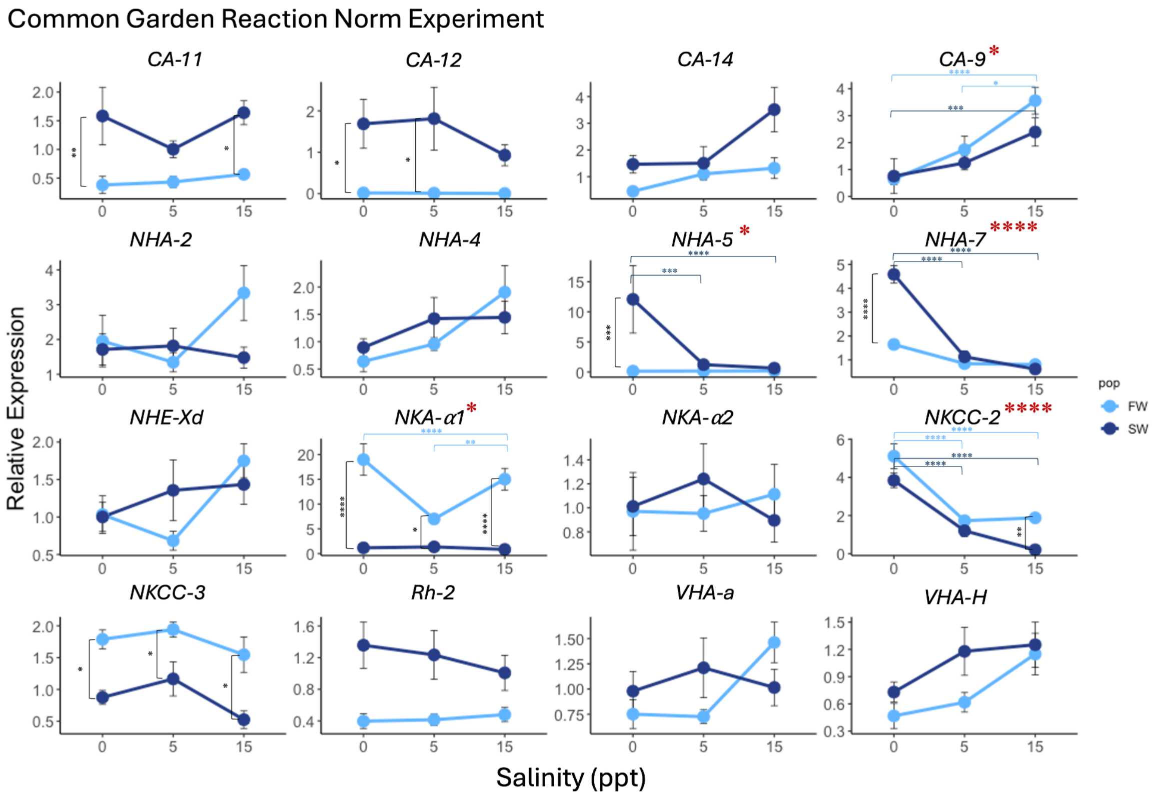


**Supplementary Figure S4. Evolutionary shifts in expression of 16 ion transporter genes between saline (SW) and freshwater (FW) populations and acclimatory shifts across the salinities (0, 5, and 15 ppt) under common garden conditions.** Relative expression refers to the amount of ion transporter (and CA) mRNA expression relative to the mean expression of housekeeping genes (*EF-1α* and *GDPH*) (Equation 1 in Methods). Data are represented as Mean ± SE across 12 biological replicates (Fig 2A). Statistical analyses were conducted using two-way ANOVA. Red * near gene name indicates a significant interaction effect between *Population* x *Salinity* (ie. evolution of plasticity) and significant results from TukeyHSD *post hoc* analysis are indicated by asterisks within each graph (*$P < 0.05$, **$P < 0.01$, ***$P < 0.001$, ****$P < 0.0001$).

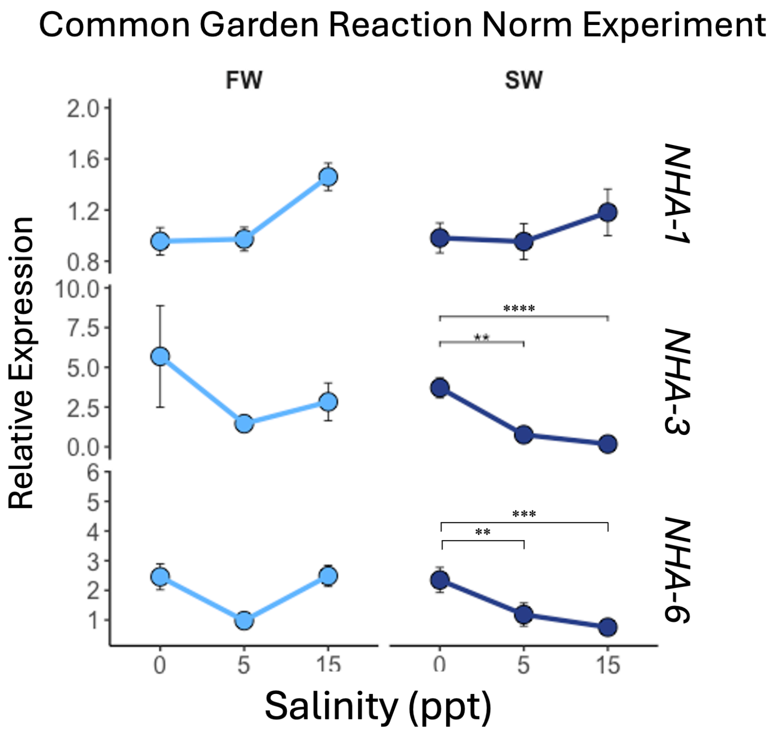


**Supplementary Figure S5. Acclimatory shifts in gene expression for NHA paralogs *NHA-1*, *NHA-3*, and *NHA-6* across the salinities (0, 5, and 15 ppt) for saline (SW) and freshwater (FW) populations.** Relative expression refers to the amount of ion transporter mRNA expression relative to the mean expression of housekeeping genes (*EF-1*$\alpha$ and *GDPH*) (Equation 1 in Methods). Data are represented as Mean ± SE across 12 biological replicates (Fig 2A). The displayed genes contained SNPs in the primer pair, and their expression is therefore not compared between the populations. Thus, gene expression across *Salinity* is compared using a one way ANOVA among the saline and freshwater population separately. Significant results from TukeyHSD *post hoc* analysis are indicated by asterisks within each graph (*$P < 0.05$, **$P < 0.01$, ***$P < 0.001$, ****$P < 0.0001$).

**Laboratory Natural Selection Experiment (Salinity decline)**

Saline population selected at 0 ppt *vs* saline population control at native salinity (15 ppt)

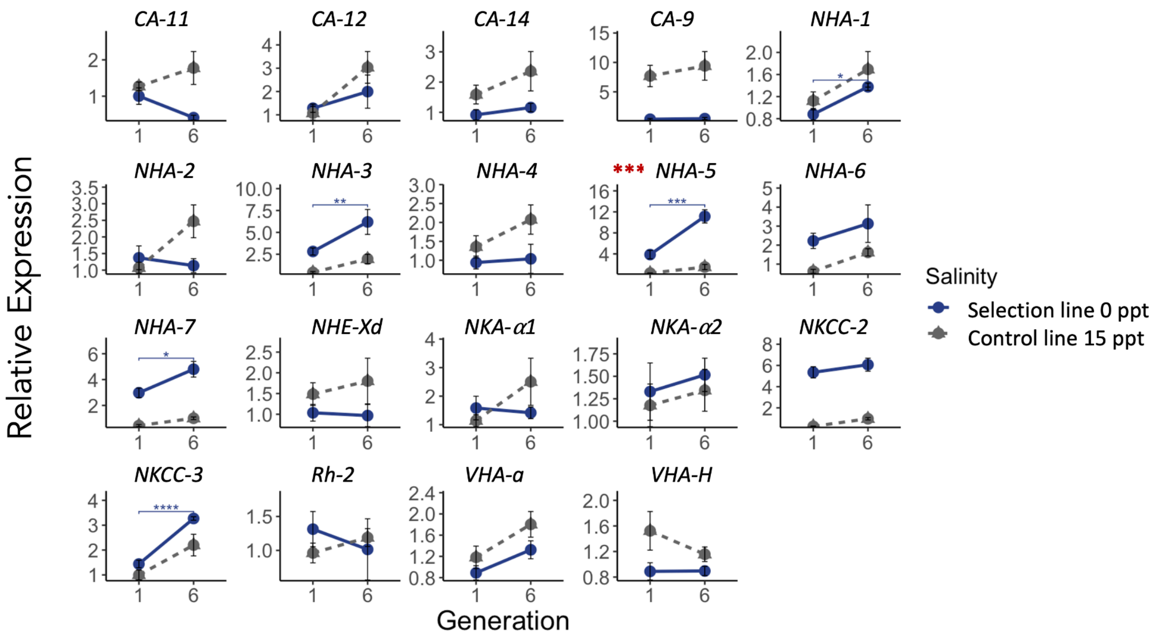


**Supplementary Figure S6 . Gene expression trajectories of iono-osmoregulatory genes in a saline population undergoing salinity decline (0 ppt) for six generations, relative to the control maintained at native salinity (15 ppt)**. Relative expression refers to the amount of ion transporter (and CA) mRNA expression relative to the mean expression of housekeeping genes (EF-1α and GDPH) (Equation 1 in Methods). Data represented as Mean ± SE. The dotted gray line is the control line maintained at native salinity (15 ppt). Red * near gene name represents an interaction effect where the slopes of the two trajectories are significantly different based on generalized linear model, navy and grey * in the graph represents significant change in gene expression within each line from generation 1 to generation 6 based on linear model. Level of significance indicated by asterisks (*$P < 0.05$, **$P < 0.01$, ***$P < 0.001$, ****$P < 0.0001$).

**Laboratory Natural Selection Experiment (Salinity decline)**

Saline population selected at 0 ppt *vs* freshwater population control at native salinity (0 ppt)

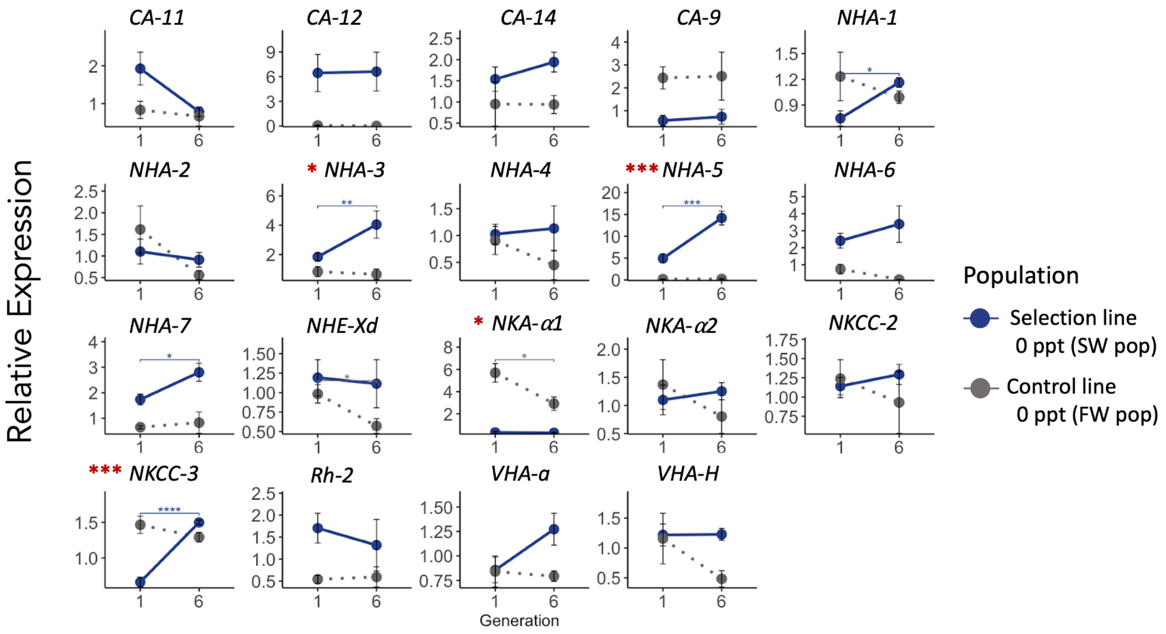


**Supplementary Figure S7. Gene expression trajectories of iono-osmoregulatory genes in a saline population undergoing salinity decline (0 ppt) for six generations, relative to the freshwater population control maintained at native salinity (0 ppt).** Relative expression refers to the amount of ion transporter (and CA) mRNA expression relative to the mean expression of housekeeping genes (EF-1α and GDPH) (Equation 1 in Methods). Data represented as Mean ± SE. The dotted gray line is the control line maintained at native salinity (0 ppt). Red * near gene name represents an interaction effect where the slopes of the two trajectories are significantly different based on generalized linear model, navy and grey * in the graph represents significant change in gene expression within each line from generation 1 to generation 6 based on linear model. Level of significance indicated by asterisks (*$P < 0.05$, **$P < 0.01$, ***$P < 0.001$, ****$P < 0.0001$).

**Laboratory Natural Selection Experiment (Salinity increase)**

Freshwater population selected at 15 ppt *vs* freshwater population control at native salinity (0 ppt)

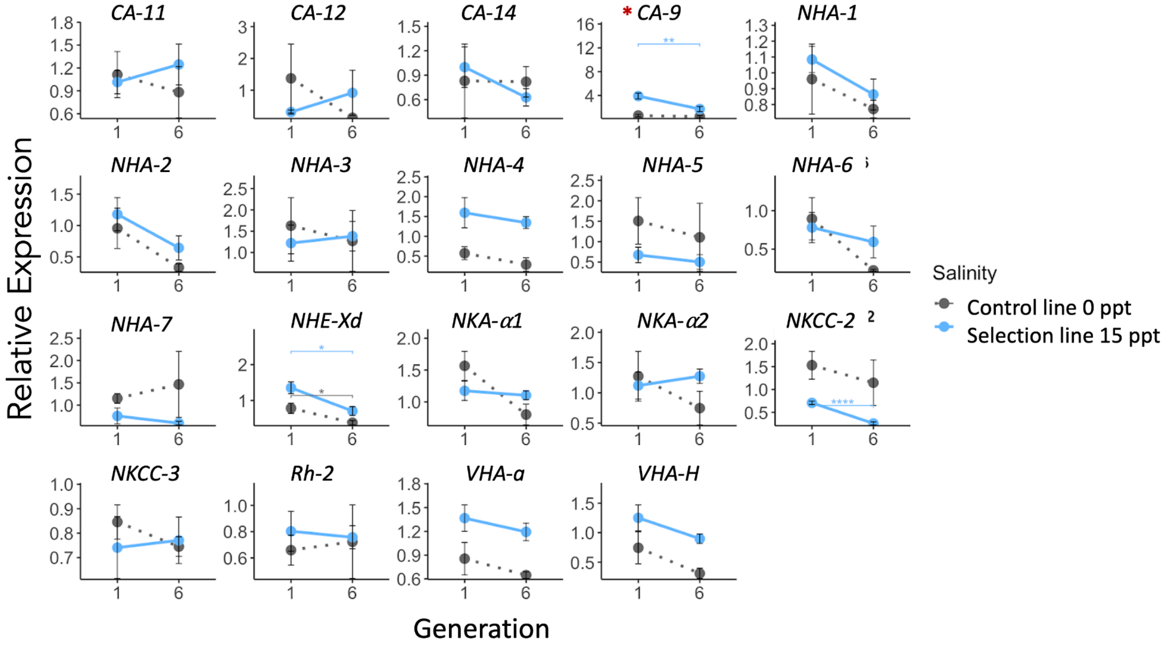


**Supplementary Figure S8. Gene expression trajectories of iono-osmoregulatory genes in a freshwater population undergoing salinity increase (15 ppt) for six generations, relative to the freshwater population control maintained at native salinity (0 ppt)**. Relative expression refers to the amount of ion transporter (and CA) mRNA expression relative to the mean expression of housekeeping genes (EF-1α and GDPH) (Equation 1 in Methods). Data represented as Mean ± SE. The dotted gray line is the control line maintained at native salinity (0 ppt). Red * near gene name represents an interaction effect where the slopes of the two trajectories are significantly different based on generalized linear model, blue and grey * in the graph represents significant change in gene expression within each line from generation 1 to generation 6 based on linear model. Level of significance indicated by asterisks (*$P < 0.05$, **$P < 0.01$, ***$P < 0.001$, ****$P < 0.0001$).

**Laboratory Natural Selection Experiment (Salinity increase)**

Freshwater population selected at 15 ppt *vs* saline population control at native salinity (15 ppt)

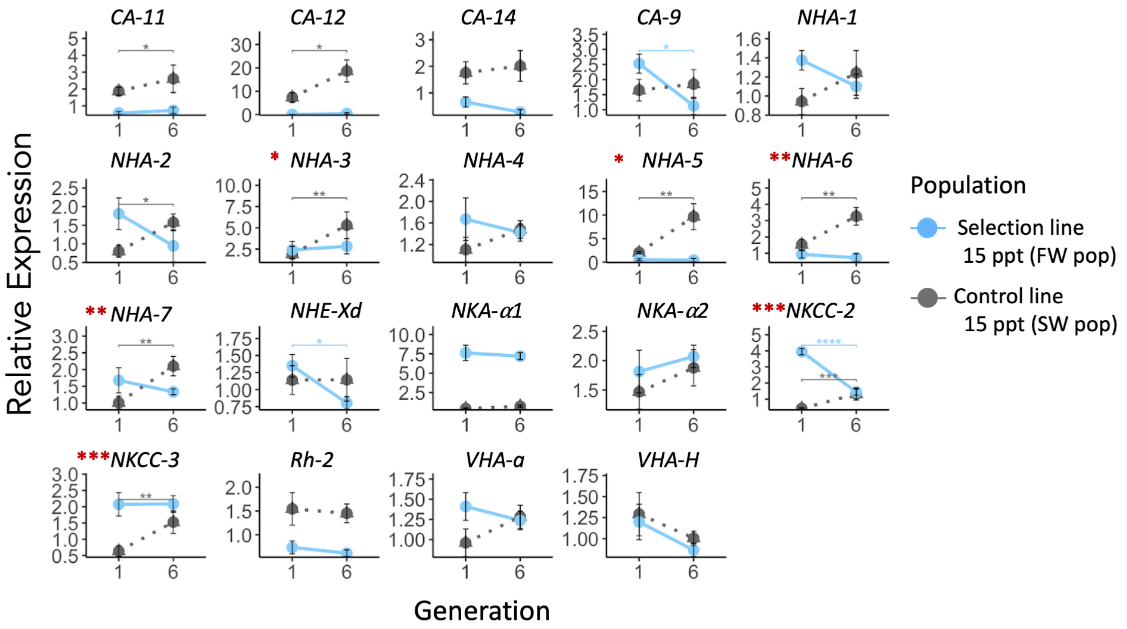


**Supplementary Figure S9. Gene expression trajectories of iono-osmoregulatory genes in a freshwater population undergoing salinity increase (15 ppt) for six generations, relative to the saline population control maintained at native salinity (15 ppt)**. Relative expression refers to the amount of ion transporter (and CA) mRNA expression relative to the mean expression of housekeeping genes (EF-1α and GDPH) (Equation 1 in Methods). Data represented as Mean ± SE. The dotted gray line is the control line maintained at its native salinity (15 ppt). Red * near gene name represents an interaction effect where the slopes of the two trajectories are significantly different based on generalized linear model, blue and grey * in the graph represents significant change in gene expression within each line from generation 1 to generation 6 based on linear model. Level of significance indicated by asterisks (*$P < 0.05$, **$P < 0.01$, ***$P < 0.001$, ****$P < 0.0001$).
